# TALC: Transcript-level Aware Long Read Correction

**DOI:** 10.1101/2020.01.10.901728

**Authors:** Lucile Broseus, Aubin Thomas, Andrew J. Oldfield, Dany Severac, Emeric Dubois, William Ritchie

## Abstract

**Motivation:** Long-read sequencing technologies are invaluable for determining complex RNA transcript architectures but are error-prone. Numerous “hybrid correction” algorithms have been developed for genomic data that correct long reads by exploiting the accuracy and depth of short reads sequenced from the same sample. These algorithms are not suited for correcting more complex transcriptome sequencing data.

**Results:** We have created a novel reference-free algorithm called TALC (Transcription Aware Long Read Correction) which models changes in RNA expression and isoform representation in a weighted De-Bruijn graph to correct long reads from transcriptome studies. We show that transcription aware correction by TALC improves the accuracy of the whole spectrum of downstream RNA-seq applications and is thus necessary for transcriptome analyses that use long read technology.

**Availability and Implementation:** TALC is implemented in C++ and available at https://gitlab.igh.cnrs.fr/lbroseus/TALC.

## INTRODUCTION

Recent advances in RNA sequencing (RNA-seq) technologies have revealed that transcription is more pervasive (1), more diverse (2) and more cryptic (3) than expected (4–6). Given the major role that RNA processing plays in disease and normal biology, it is crucial to ascertain the existence of novel isoforms and to accurately quantify their abundance. Second generation RNA-seq technologies such as Illumina are well suited to the tasks of assessing gene expression levels and determining proximal exon connectivity. They produce numerous sequencing reads at a low cost ensuring sufficient representation of most transcripts. However, because the RNA or cDNA is fragmented during short RNA-seq protocols, long range connectivity can only be computationally inferred. These predictions based on short reads struggle to correctly identify transcript isoforms that contain multiple alternative exons (7) or that contain retained introns (8, 9). In these cases, long-read (LR) sequencing technologies are invaluable because they can sequence entire molecules in one pass and thus capture long-range connectivity of complex isoforms.

LR technologies however produce less reads than short read (SR) sequencing approaches (10) for similar costs and have higher error rates. In many cases these higher error rates can prevent the correct identification of isoforms (11–13). Although several alignment software (14–18) are optimized to handle these errors, their shortcomings confound transcript identification and annotation. Many reads cannot be aligned and regions where the sequencing error rates are higher such as UTRs frequently produce ambiguous alignments. Secondly, they struggle to identify splice junctions, notably those flanking small structural variants such as small exons or alternative 5’ and 3’ splice sites. This impacts the evaluation of exon skipping events and worsens the quality of transcript assembly (12). These drawbacks prompted the development of algorithms, referred to as *hybrid methods* (19–22), that take advantage of SR depth and accuracy to compensate LR shortcomings.

Numerous algorithms have been proposed to combine long and short reads into high-accuracy long reads (23). A first approach is to correct long reads by local consensus inferred from short read multi-alignments (24–27). This strategy is generally slow and computationally intensive. More importantly, it tends to show poor performances over low-expressed regions, and a risk of bias toward major isoforms. Therefore, they may not be suitable for transcriptome and single cell datasets where isoform representation may vary considerably. In the second approach (28–31), short reads are considered as the fragments of a reference transcriptome, and roughly assembled using a graph structure, onto which long reads are aligned (32). However, sequencing errors in SR datasets, complex transcript architectures and sequence duplicates often lead to extremely complex graph structures (33, 34) that elicit graph simplifications or exploration heuristics to keep computations tractable. Most hybrid correction algorithms were primarily intended to be applied to genomic data with the aim to improve the quality of genome assemblies. As such, they typically rely on assumptions that fit DNA-seq data properties and thus a linear reference genome. According to this, bifurcation nodes and tips (dead-ends in the graph) are assumed to originate mainly from sequencing errors and not transcript processing events. In addition, genome sequencing benefits from a relatively uniform read coverage and therefore the graph is simplified by discarding *a priori* all nodes whose count is below a user-defined threshold. The fact that these approaches are not adapted for RNA-seq data have been extensively discussed in previous works on transcript assembly (34) and short read correction (35, 36) and their impact on long read correction can be easily anticipated. RNA-seq data display highly uneven coverage, even across a same genomic location, thus the frequency of sequencing errors varies depending on the surrounding coverage depth. Tips may correspond to the start or end of a transcript and finally specific regions such as 3’ ends of transcripts, are frequently under covered.

We have developed TALC for Transcription Aware long-read Correction which addresses RNA-seq data specificities by incorporating coverage analysis throughout the correction process. TALC considers transcript expression and the existence of isoforms to correct LRs. For the purpose of testing TALC, we generated Oxford Nanopore direct RNA sequencing reads and Illumina short reads on an MCF10A human cell line and downloaded LR and SR data from the GM12878 human cell line and a SIRV Spike-In experiment. We demonstrate that after TALC correction, long reads map with higher sequence identity and with less errors in exon assembly than currently used methods. The gains observed following TALC were tested on simulated long reads as well as on real reads.

## SYSTEM AND METHODS

### Methods overview

Figure 1 illustrates the methodology behind TALC and highlights how it considers transcript abundance and architecture to correct long reads. The first step of TALC graph-based procedure is to consider short reads as a raw reference transcriptome by merging them through a *weighted De-Bruijn Graph* (DBG) structure (28, 32), whose nodes represent k-mers. For each node, we record their k-mer abundance in the SR dataset. Thus, any sequence of transcripts expressed in the RNA-seq sample should appear as a unique path of the graph. The long read is corrected by finding the right sequence of nodes to which it corresponds in the *DBG* built from short reads. Paths corresponding to true transcripts should display consistent k-mer coverage except in regions where the existence of multiple isoforms may alter the coverage such as at splice junctions (Figure 2). We thus propose a method of graph exploration that considers k-mer count variation consistent with transcript abundance and isoform existence (adaptive count thresholding).

**Figure 1:**
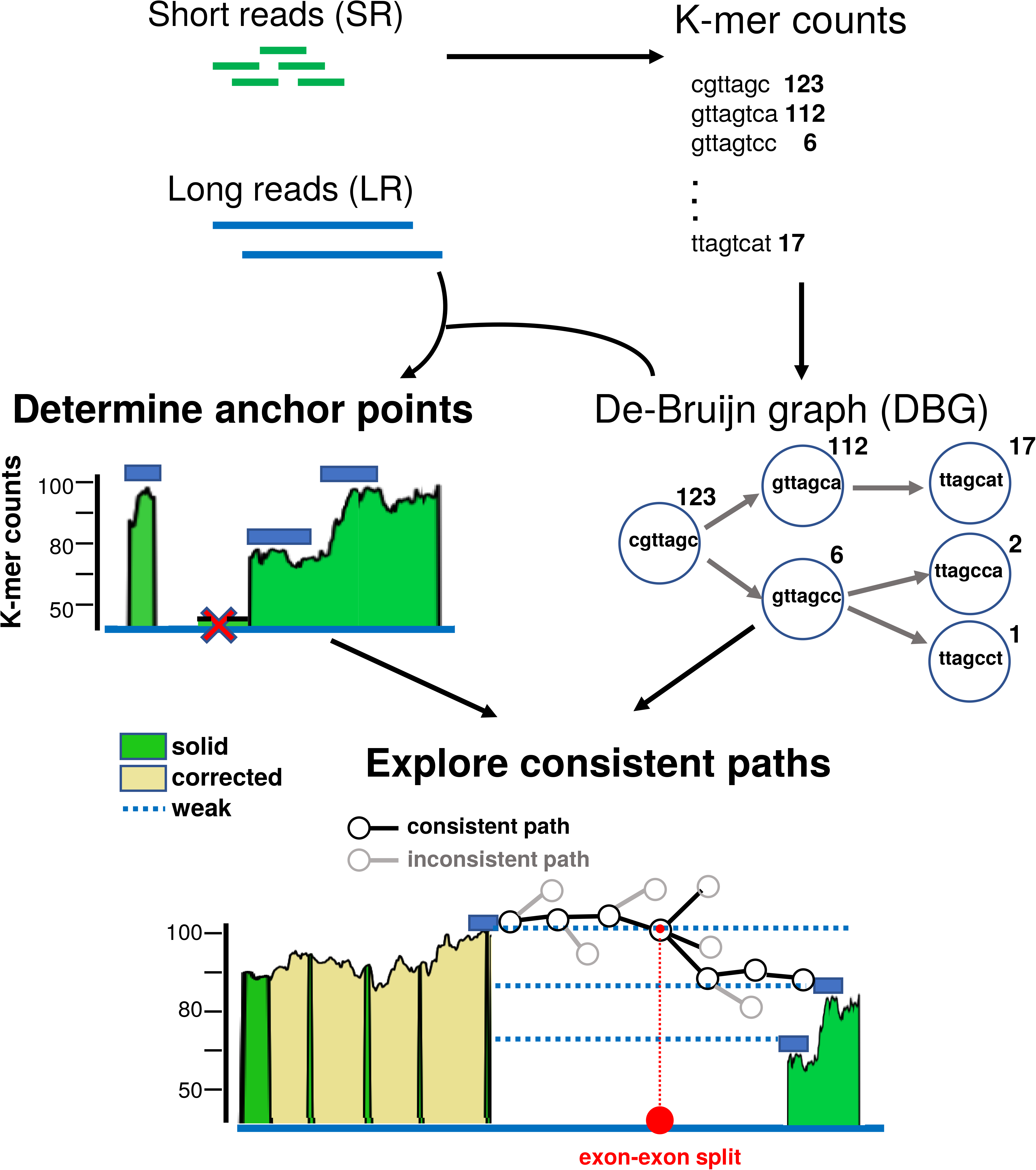
Overview of the TALC algorithm. TALC’s correction procedure begins by creating a weighted DBG (DBG) from the k-mer counts of the SRs. It then searches for common sequences between the LRs and the k-mers of the DBG. Stretches of common k-mers are called *Solid Regions* and are assumed to be error-free parts of the long read (green peaks). From the solid regions, TALC will then **determine anchor points**. These will be used as entry points into the DBG; regions between these anchor points are called *weak regions* and will be corrected. To extract anchor points from the solid regions TALC will first crop background noise that frequently surround solid regions (red cross). The second step in determining anchors is to search for sudden changes in coverage within a solid region. These changes may correspond to divergent transcript architectures and should be explored separately. Thus, a solid region will be split into as many anchors as there are changes in coverage. Finally, to correct weak paths between anchors, TALC will **explore consistent paths.** TALC explores the DBG, following paths of similar k-mer coverage. When the exploration reaches a potential transcriptional event such as a splice site (red dot), the exploration will branch out to account for the existence of multiple isoforms with varying coverage.

**Figure 2:**
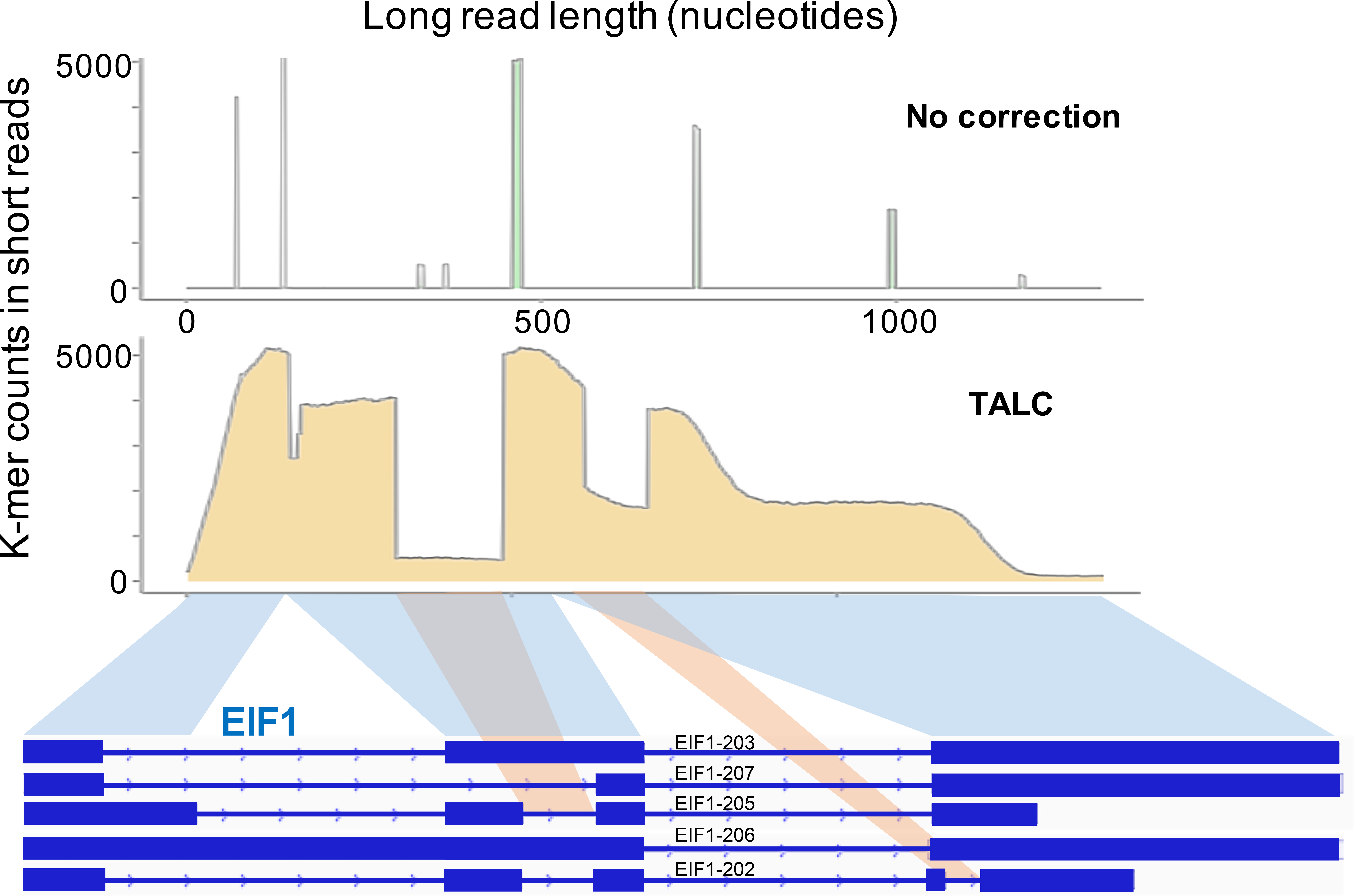
Example of K-mer coverage across a long read before and after TALC correction. We mapped the k-mers discovered in the short-read (SR) sequencing of MCF10A cells onto one of the long reads (LR) sequenced from the same sample. The x-axis represents each nucleotide of the LR, the y-axis shows the abundance of the mapped k-mers in the SR sample. Blue shading at the bottom of the figure highlights the correspondence between the transcript to which the long read was mapped and the read itself; red shading highlights alternative transcript architectures that could explain the sudden changes in the k-mer coverage of the graph. Alternative isoforms are annotated using ENSEMBL transcript annotation.

### Determining anchor points

As in (28), the LR sequence is first divided into *solid regions* (stretches of k-mers shared with the SRs) interspersed with w*eak regions* (stretches of likely erroneous k-mers).

To crop background noise that frequently surround solid regions, we first estimate the frequency of k-mers resulting from sequencing errors in the SR. The count of k-mers containing a sequencing error in SR is assumed to follow a Poisson distribution whose mean should be less than the average noise level λ. An estimate of λ is taken as the average coverage depth across the LR multiplied by the per base error rate in SR, whose estimates were found to be higher in RNA-seq datasets than in genomic data, ranging from 1% to 3% (35).

A robust estimate of the average depth is derived *a priori* from the shared k-mer counts: the 15% smallest and 10% highest counts are removed and the mean is calculated from all the remaining k-mers. K-mers whose count falls under the upper bound of a confidence interval from the Poisson distribution Poisson(λ) (cf: Supplementary Materials) are not considered solid and will not be used as anchor points.

We then split solid regions into anchor points with contrasting k-mer coverage. To this end, we inspect the regions coverage and each k-mer at which an abrupt count variation is detected is selected (cf Competing paths section below). K-mers at the tips of a region are always selected, and additional k-mers may be picked so as to have at least three distinct anchor points per solid region.

Once the long read is anchored, the DBG is explored in-between any couple of consecutive anchors, in search for the error-free path matching the inner weak region.

### Selective Exploration of the DBG

In TALC, a path in the DBG is defined as an ordered list of connected k-mers (nodes) which are weighted by their number of occurrences in the SR dataset. When a k-mer is mapped to a LR, we attribute its weight to the mapped region of the LR and use the term coverage. Thus, when we speak of coverage of a LR, it is in effect the weight of the k-mer that mapped to that region.

Abrupt changes in the coverage depth can be caused by alternative splicing events, duplicate k-mers, or sequencing bias in SR. These abrupt changes are termed split-coverage.

Nodes in a *DBG* are either *simple,* which means there exists only one successor k-mer in the data, or admit at least two successors, and thus can be extended by several distinct paths. We will refer to them as *bifurcation nodes*. Exploration is implemented as a breadth-first approach where at each bifurcation, only the next nodes whose count is considered consistent with the coverage of the current path will be followed. More precisely, at a given step n0, the k-mers database is queried for the counts of its four possible successors: (nA, nC, nG, nT). And we want to assess which of these make up valid extensions of the current path.

We need to infer two things: whether n0 is close to a split-coverage event, in which case the exploration might be split into several competing paths; and which nodes are likely erroneous and should be filtered-out. To this end, we use the following decision rules (cf: Supplementary Materials):

1. If there is only one non-zero k-mer, there is no decision to make.
2. If there are at least two non-zero k-mers, we postulate that most of the time, there is no real split coverage event at n0, so that its count provides a good estimate of the downstream coverage depth, from which both the correct successor (say nL) and a local noise threshold can be inferred.

Accordingly, allowing to the consistence of the coverage, we pose as a null model:

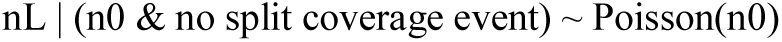

and

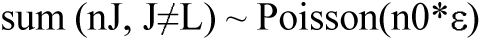

When the counts (nA, nC, nG, nT) do not fit this null model, we infer the current bifurcation is due to a change-point (eg: an exon junction site with several splicing variants) and the exploration is split into all downstream nodes. At each new node, the expected count and the noise level are thus re-estimated, allowing an adaptive filtering-out of erroneous k-mers.

Therefore, a true path is expected to be mostly made of count-consistent nodes possibly with a few change-point nodes. And the more a path admits change-points the less likely it is to represent the sequence of a true transcript.

### Competing paths

TALC favours coverage-consistent exploration of the DBG. As opposed to LoRDEC (28), all paths passing the test described above are explored in parallel by a breadth-first approach. When the number of parallel paths exceeds a specified threshold, an evaluation is performed: all on-going paths are compared to the LR sequence and the least similar ones are stopped. This allows a more local and gradual evaluation of the similarity between the paths and the long read and we believe it contributes to reduces the inclusion of small sequencing errors from the SRs into the LR.

All paths which successfully bridge both Solid Regions are scored by their edit distance with the LR. The most similar one is considered suitable for further validation. Its sequence is aligned against the LR’s; if the percent identity score with the LR is higher than a user-specified threshold (by default 70%), we assume we have likely found the best candidate.

### Border exploration

Compared to inner weak regions (flanked on both sides by a solid region), the correction of weak border regions (flanked only on one side by a solid region) raises additional problems. Firstly, there is no clear targeted anchor point at which to stop exploration. Secondly, UTR sequences often contain low complexity regions and duplicated k-mers, which leads to increased complexity in the DBG. Finally, borders of transcripts suffer from higher sequencing bias in short reads (notably in 3’ ends of transcripts). The two last points make k-mer coverage more erratic and over dispersed. In certain cases, incomplete coverage UTR extremities (eg: rare longer UTR forms) can sometime prevent a full-length correction entirely. For these reasons, we rely more heavily on sequence length and similarity between the visited paths and the LR’s to direct graph exploration across those regions. More exactly, graph exploration is monitored in a same manner as described above (cf: **Selective Exploration of the DBG)** until the paths’s length matches the border’s length. When the number of consecutive errors exceeds a given threshold, the corresponding paths are stopped.

When there are no more branches to investigate (that is all possibilities have ended in dead-ends of the graph or have been stopped), the very last error-prone bases of all interrupted paths are trimmed (cf: Supplementary Materials). To elect the best border path, we first search for it among the paths that were at least as long as the portion of the read we are attempting to correct, despite the constraints on sequence similarity. We choose the one having the smallest edit distance with the LR, and the most consistent coverage if there are *ex-aequo*. If no long border path could be found, shorter paths are compared and the one with highest similarity to the LR is selected.

### Choosing between multiple paths whose scores are tied

Paths in the *DBG* that successfully bridge two Solid Regions are first ranked according to their sequence similarity with the Weak Region’s. The similarity is computed as the edit distance between sequences.

We notice that the exploration stage often provides several solution paths having the exact same alignment score (this occurs for example when there are variants in sub-sequences that have been inserted or deleted from the LR), so that we cannot decide between them based on sequence information only. In order to choose between these equally-scoring paths, we perform a second ranking using count variation: the path with the least change-points wins.

## EVALUATION AND RESULTS

### Benchmarked algorithms

We compared the corrections made by TALC with six other state of the art hybrid correctors: FMLRC (31), CoLoRMap (26), Jabba (29), Hercules (27), LoRDEC (28) and LSC (24). We chose FMLRC, Jabba, LoRDEC and ColoRMap as they were found to be the top performers according to a recent benchmark on genomic data (37). We added Hercules because it was a very recent method not yet benchmarked. We also included LSC for it is (with LoRDEC) currently applied in numerous transcriptomic studies combining long and short reads (38–41).

Though it was well suited for RNA-seq data, according to (42), we could not include NaS (43) as it depends on a third-party proprietary software (Newbler assembler) that is no longer available.

Several algorithms (eg: CoLoRMap, LSC and Jabba) exclude uncorrected reads from the final output file. However, to allow a fair comparison with the other methods, we systematically added back all discarded raw sequences before evaluation.

All jobs were run using 15 CPUs with 512 gigabytes of available RAM. If a job was not finished after 1 week, it was terminated (Jabba failed to finish within a week and had to be finally remove from the analysis; as already observed in (27), LSC could not scale to the biggest size real LR datasets; Hercules is absent from several tests too, due to long running times and heavy computational requirements, chiefly in its short read alignment phase). LoRDEC and TALC were run with the recommended size of k-mers set to 21 (28, 44), and FMLRC with its default parameters.

### Benchmark datasets

To evaluate the efficiency of these algorithms, we made use of two publicly available and one in-house SR and LR matched real datasets. The first dataset comes from a large GridION MinION cDNA sequencing experiment of SIRV E0 Spike-Ins (45). The second one is provided by the Nanopore consortium (4) and was generated on the GM12878 B-Lymphocyte cell line. Because there was no SR dataset provided with this experiment, we used the GM12878 Illumina data from an a earlier study (cf: Supplementary Material). Lastly, we performed our own Oxford Nanopore (ONT) direct RNA sequencing and matched Illumina mRNA sequencing on the MCF10A cell line (cf: Supplementary Material).

By using real datasets with contrasting error models (cf: Supplementary Material) to compare these methods instead of a computationally generated one we are testing them in authentic conditions. However, the drawback of using real data is that it is impossible to measure precisely the number of good corrections performed because the entire repertoire of transcripts and their true abundance is unknown; there is no reference transcriptome that perfectly matches the MCF10A and GM12878 cells. For example, (4) reconstructed about 78 000 high-confidence isoforms from the GM12878 datasets and found that the majority were absent from reference databases (4, 46).

To work around this problem, we used the (raw) LR data to measure the abundance of transcripts in MCF10A and GM12878 cells (Supplementary Materials). From this LR-derived transcriptome, we simulated long ONT reads that respect the error rates that we observed in the sequencing data. Thus, we have a dataset of reads that are derived from the MCF10A and GM12878 transcriptomes and for which we know the exact structure and abundance of each isoform. This enables us to evaluate performances on a simulated LR dataset derived from the MCF10A and GM12878 transcriptomes.

### Measures of evaluation

We wished to use the recently published evaluation tool, *LC_EC_analyser (42)*. Unfortunately, it does not scale to our real size datasets (seemingly because of AlignQC issues in memory management). Nonetheless, we used analogous criteria (see below). Scripts designed to perform our evaluation are avalaible at: https://gitlab.igh.cnrs.fr/lbroseus/TransAT.

To assess the behaviour and performance of the 7 correction methods, we computed multiple indicators that are summarized in Radar Charts and Supplementary tables 1-3. These indicators can be broken down into two categories: *standard sequence quality* indicators and *transcriptome-specific* indicators.

#### Standard sequence quality indicators

These include the base accuracy between the LR and the molecule from which the LR was generated, the various error rates (mismatches, insertions, deletions), and the percentage of primary alignments to the reference genome. They are common benchmarks for genomic data. In addition, we verified that the different correction algorithms were able to maintained the initial read length.

#### Transcriptome-specific indicators

When evaluating correction methods on real data, performance metrics can be unintentionally skewed because the molecule that was sequenced is unknown. For example, a correction method that would simply replace a LR by a known transcript sequence to which it was most similar could obtain a high percentage of mapped reads and low error rates but would not correctly represent the sequenced transcriptome. Thus, to further evaluate the 7 correction methods in this paper, we computed two sets of transcriptome-specific indicators which are directed towards the two major applications of RNA-seq experiments: transcript-level quantification (11, 47) and isoform recovery (12). For this we considered two types of data: a real SIRV Spike-In dataset and two simulated LR data mimicking the real MCF10A and GM12878 experiments. SIRV Spike-In data have known theoretical concentrations from which we could gauge how each correction method would impact transcript counts accuracy. Simulated LRs datasets mimicking the real MCF10A and GM12878 datasets both in terms of error-model and transcript-level expression (cf: Supplementary Material) allows us to extend our observations on SIRV data to more complex transcriptomes. Importantly, it gives us insight into the capacity of a hybrid corrector to preserve and clarify transcript structure at the read resolution. For measuring the impact of correction on structure and transcript assembly accuracies, we computed the proportion of (simulated) long reads whose transcript structure could be correctly elucidated by aligners and their ability to preserve true exon connectivity and to improve false ones (cf: Supplementary Tables 2C and 2D).

### Sequence quality of long reads

We measured the general quality of corrected sequences using the percentage of mapped reads, the base accuracy and the type of errors that remained in the sequences. These are shown in Figure 4. Our first measure of performance was the number of mapped reads following correction (Figure 4, Supplementary Table 1A). This was evaluated for all 7 methods using three different aligners: Minimap2 (15), GMAP (14) and GraphMap (16). For all three aligners, we kept default parameters suggested for ONT data by their respective authors. We chose Minimap2 and GraphMap as they were specifically designed for long mRNA reads and GMAP because it was considered the best splice-aware aligner for long RNA data according to a recent benchmark (48). TALC shows consistently high mapping numbers regardless of the aligner used. It’s closest competitor in this aspect, LoRDEC, has a high number of mapped reads with Minimap2 and GMAP but this drops to the lowest number of all methods with GraphMap. We noticed a similar trend at the gene-level assessment where LoRDEC correctly associated LRs with their gene of origin when Minimap2 or GMAP were used but was one of the worst performers with GraphMap (Figure 4, Supplementary Table 2A). This can be explained by the rather high mismatch rate that remains after LoRDEC correction (partly due to the insertion of false SNPs from SRs sequencing errors, Supplementary Table 1D).

Based on these alignments, we computed the *base accuracy* of corrected reads as in (47) (Figure 4, Supplementary tables 1B to 1D). To assess whether the benchmark performances depended on the coverage in SRs, we also computed the summary statistics of base accuracy according to the gene coverage depth in SRs (Supplementary table 1E). The results of this analysis show that TALC globally performs well on this measure and across the entire range of SR coverage bins. It’s closest competitors for base accuracy are CoLoRMap and FMLRC.

### Read length conservation

Long read technologies are often used to determine entire transcript architecture including alternative start or end sites (49). Hence, it is important that correction algorithms include a strategy to delimitate RNA read borders accurately. To compare the length of corrected sequences to the raw read, we calculated the relative distances between the raw and the corrected read length over all real datasets (cf: Supplementary Table and Supplementary Figure 1F). Overall, the methods considered seem to control read length with an average 2% to 3% difference in length with the raw read. CoLoRMap was the only exception as we found that it extended reads with an additional 5%-13.61% which could cause the correction to drastically overextend the transcript borders.

### Exon structure preservation and clarification

LR technologies are able to capture the full connectivity between exons of transcripts. Although this does not necessarily require nucleotide resolution of the transcript, the error rate of current LR technologies is sufficiently high to confound exon connectivity analysis (Figure 3A). Here we define “structural errors” as the incorrect inclusion or deletion of an exon or the incorrect identification of exonic boundaries. We developed a method to systematically identify these errors (Figure 3B and Supplementary Material) and evaluated the impact of LR correction methods on them. Because the mapping approach used to assign LRs to a given transcript may impact on structural errors we again tested three mapping approaches: mapping of LRs to a reference genome (hg38) using GMAP (14) and Minimap2 (15), mapping of LRs to a reference transcriptome (hg38, ENSEMBL release 97) using GraphMap (16). In our evaluation, we measured for each exon, how many were properly identified after correction (Supplementary Tables 2C and 2D). If these exons were already properly identified before correction, we say that they were preserved by the correction algorithm; if they were not, we say that they were clarified by the correction algorithm (Supplementary Table 2D). These results are summarised in Figure 4B.

**Figure 3:**
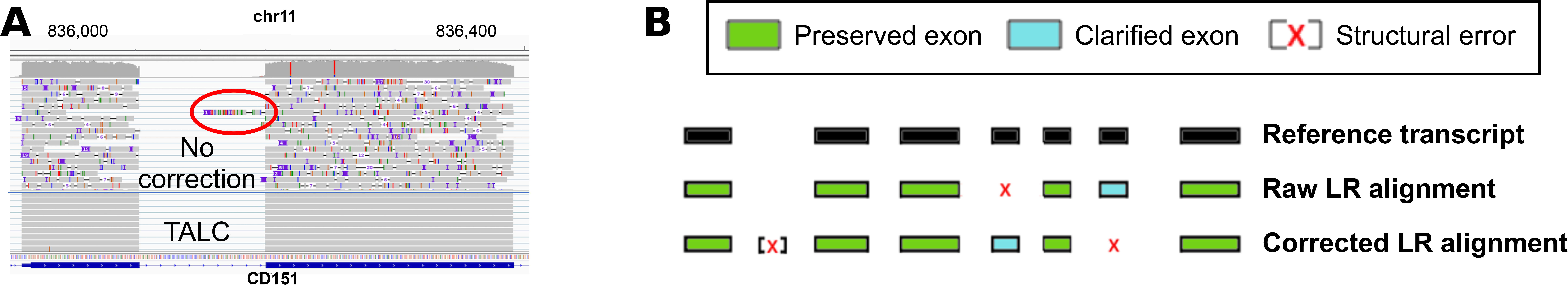
Structural errors in long reads after different correction methods. **A)** Screenshot of the IGV viewer showing long reads aligned to the genome with GMAP. A structural error (insertion) has occurred in the non-corrected long reads (red circle). **B)** Overview of our approach to finding structural errors which are either deletions or insertions in LRs computationally generated from a given transcript.

**Figure 4:**
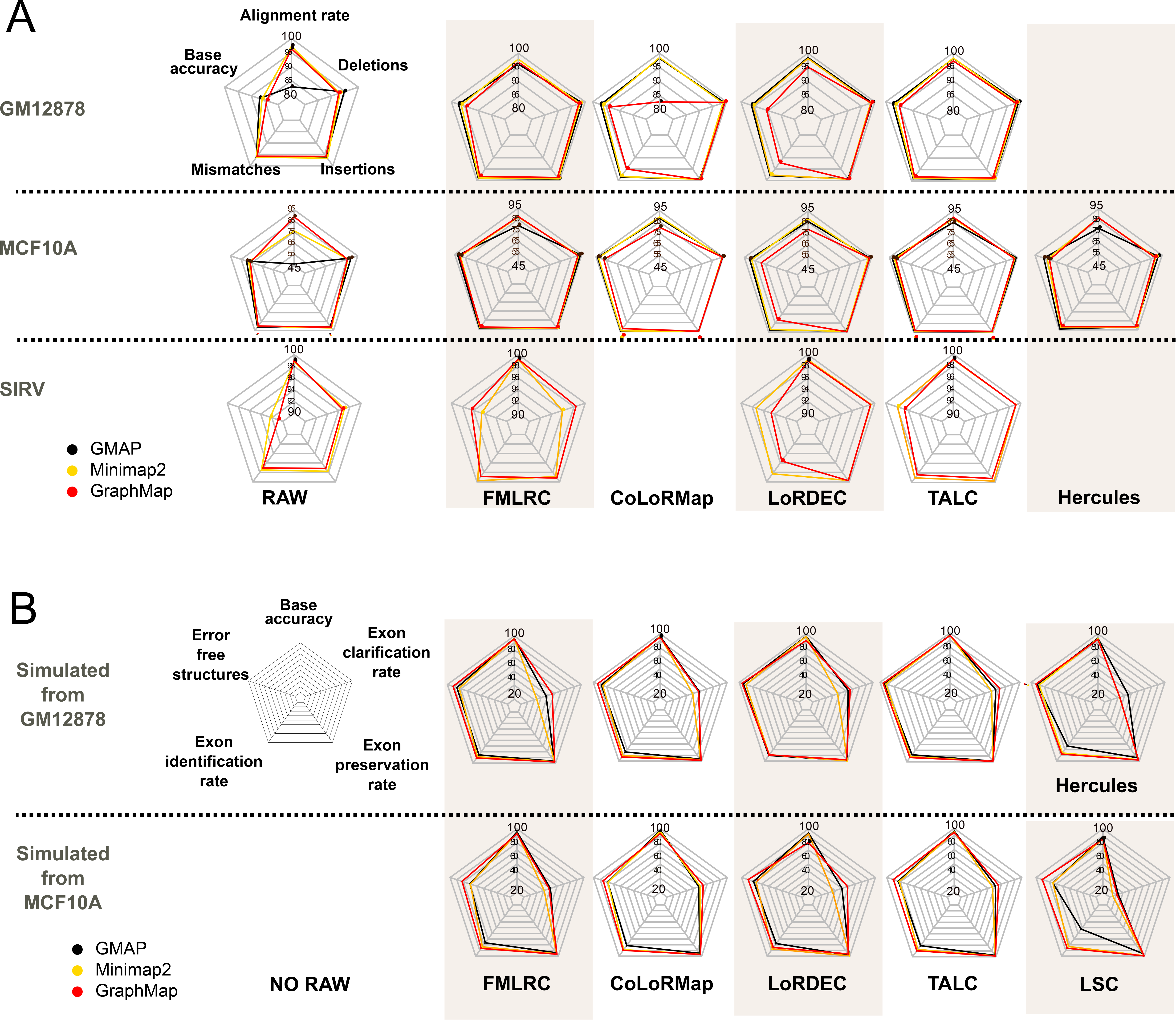
Benchmark of correction algorithms using sequence and transcript features. **A)** Radard charts of sequence specific measures of correction efficiency for cell lines GM12878 and MCF10A and for the SIRV spike-in set. **B)** Radard charts of transcript structure-specific measures of correction efficiency for data simulated from cell lines GM12878 and MCF10A. Missing plots for CoLoRMap, Hercules and LSC mean that these algorithms were incapable of running properly on the given dataset.

Regardless of the mapping strategy, we found that TALC is systematically in the top two best algorithms and globally performs the best (Figure 4B). Again, we notice that other algorithms may compete with TALC on specific criteria given specific aligners but their performance drops drastically in other conditions. For example, when using the Minimap2 aligner, LSC slightly outperforms TALC in preserving the number of properly assigned exons (99.3% versus 98.5% preserved exons) but is very poor at correcting exons that were initially incorrect (14.8% versus 56.6% clarified exons).

### Gene and Transcript-level quantitation

To our knowledge, at the time, no algorithm dedicated to long read transcript-level quantitation has been published nor validated yet (11, 47). The authors of (47) tested several strategies to assess whether Nanopore data were fit for quantification. We reproduced their approaches based on Salmon (Salmon *quasi-mapping*). When splice-aware genome alignment was applicable, we also tested the approach *Minimap2+Salmon.* For comparison, we added a third approach, independent of Salmon, and simply based on GraphMap transcript assignments. Quantitation was realised on real SIRV Spike-In data (transcript-level) and on simulated datasets (gene-level and transcript-level). On simulated data, this analysis suggests that read correction results in a significant overall improvement of gene-level quantification accuracy (cf: Supplementary Table 2A), all correction methods providing a rather similar gain in accuracy. This observation seems to hold also at the transcript-level, when using Salmon quasi-mapping mode, both on simulated datasets (Supplementary Table 2B) and real Spike-In data (cf: Supplementary Table 2E). But, surprisingly, a prior alignment step (using either Minimap2 or GraphMap) seem to level the estimates.

## DISCUSSION

High error rates of long read sequencing technologies can substantially bias the assignment of reads to unique transcripts and can also introduce major structural errors in *de novo* transcript prediction. Consistent with a previous study (12), we show that proper hybrid correction can provide significant improvement in the quality of downstream transcriptome analyses. However, existing hybrid correction algorithms were designed originally to improve genome assembly and they are not all suitable for RNA-seq data (42).

We propose a hybrid correction method tailored to RNA sequencing that considers transcript abundance to detect the possible existence of splice junctions and correct RNA long reads. This information is used to guide the exploration of a graph structure by eliminating edges that are not consistent in terms of transcript abundance while simultaneously using an adaptive threshold to account for the existence of multiple transcript isoforms. By eliminating inconsistent nodes, TALC reduces the inclusion of false nucleotide variants present in short read data. And by integrating coverage information, it can efficiently detect and account for the existence of multiple transcript isoforms.

We tested the efficiency of our approach on three real and two simulated datasets. Globally, TALC shows better correction than currently used methods both in base accuracy and alignment rates but more importantly it conserves the exon connectivity in transcripts. Although other state of the art correction algorithms compete with TALC on specific applications, they have clear deficiencies on other important criteria that we summarize here.

**FMLRC** was the fastest software included in our benchmark. Additionally, it does very well on base accuracy. However, it showed poor performances over most transcript-specific measures. This behaviour may be due to its approach for filtering “erroneous” k-mers. The threshold is read-specific but is computed *a priori* from the median SR read coverage (31). This likely deletes minor isoforms k-mers when the difference in coverage with the major isoform is high. In addition, we noticed that FMLRC tended to trim reads.

Although it has long running times on real size datasets (cf: Supplementary Table 3), **CoLoRMap** exhibits interesting properties when applied to RNA-seq data. Analysis on simulated data indicates it can fairly improve the reconstruction of the inner body of transcrits. Nonetheless, it lacks a specific strategy to stop correction at the borders, and has a marked tendency to extend reads. This feature, though relevant for genome assembly, can be detrimental on RNA-seq data when the corrected read originates from a gene which expressed multiple UTR forms or undergoes alternative transcription start and end.

Consistent with conclusions from (42), **LoRDEC** demonstrates very good results on most the transcriptome-specific indicators. Its main drawback is a lower base accuracy due to the inclusion of many sequencing errors from the SRs into the LRs, especially across high SR coverage regions. Additionally, it’s performances vary dramatically between aligners.

**TALC** displayed a good capacity to improve sequence quality while preserving splicing variants and proved robust to all error profiles considered. In addition, it provided rather consistent results whatever the aligner that was used across all aligners. Overall, TALC is systematically amongst the best performers across all metrics seems fitted to improveand improves the quality of transcriptome assemblies.

Given the recent discovery of numerous novel functional transcripts in multiple organs (3, 58), the re-evaluation of transcript diversity and complexity in model organisms (4, 5, 59, 60) and the growing interest in Oxford Nanopore direct-RNA multiple assets in viral transcriptome research (61–64), TALC’s capacity to correct the entire gamut of transcripts, and preserve the correct transcript structure may prove increasingly valuable.

## DATA AVAILABILITY AND IMPLEMENTATION

TALC is written in C++ and uses the SeqAn library (65, 66). The program is freely available under the CECILL licence in the GitLab repository (https://gitlab.igh.cnrs.fr/lbroseus/TALC). Currently, the De Bruijn graph is built from k-mer counts files as output by Jellyfish2 (67).

All R scripts written to evaluate hybrid correction methods can be found at https://gitlab.igh.cnrs.fr/lbroseus/TransAT.

Direct RNA Nanopore and Illumina RNA-seq MCF10A samples have been deposited on GEO under accession number GSE126638.

## AUTHORS’ CONTRIBUTIONS

LB developed and implemented the algorithm. LB and WR devised the analyses and planned the experiments. AT provided valuable suggestions for the implementation. AJO cultured MCF10A cells and extracted their RNA. DS carried out the sequencing, and ED performed the base calling of the MiNION sample. LB and WR wrote the article. All authors read and approved the final manuscript.

## ACKNOWLEDGEMENT

We wish to acknowledge the Genotoul platform (genotoul.fr) for providing us with calculation time on their servers.

## FUNDING

This work was supported by the Agence Nationale de la Recherche [ANRJCJC - WIRED], the Labex EpiGenMed [ANR-10-LABX-12-01] and the MUSE initiative [GECKO].

DS acknowledges financial support from France Génomique National infrastructure, funded as part of “Investissement d’avenir” program managed by Agence Nationale pour la Recherche [ANR-10-INBS-09].

## CONFLICT OF INTEREST

The authors declare that they have no competing interests.

## Notes

https://gitlab.igh.cnrs.fr/lbroseus/TALC

### Competing Interest Statement

The authors have declared no competing interest.

### Summary of Updates

Section Evaluation has been updated to include new results on SIRV Spike-In data and new figures. The supplemental file has been updated accordingly.

