## Supplementaty Materials for "TALC: Transcript-level Aware Long Read Correction"

***RNA sequencing with Oxford Nanopore Technologies***

Poly(A)+ RNA is prepared from 75 µg of total RNA using the Dynabeads mRNA Purification kit ref 61006 (Thermofisher). This process purifies the poly-A containing mRNA molecules using poly-T oligo attached magnetic beads using two rounds of purification. For the second elution mRNA is eluted in 20µl of pure water. The quality and the quantities of mRNA are checked on the Fragment Analyser with HS RNA sensitivity kit (Agilent).

Direct RNA libraries were prepared with the Direct RNA kit SQK RNA01 from Oxford Nanopore Technologies. 500-1280ng of poly(A)+ RNA was ligated to a poly(T) adaptor using T4 DNA ligase. The ligated RNA was copied into first strand cDNA using superscript III reverse transcriptase. The products were purified by adding a 1.8-fold excess of Agencourt RNAClean XP beads and following the Agencourt purification protocol. Sequencing adaptors preloaded with motor protein were then ligated onto the overhang of the previous adaptor using T4 DNA ligase. Excess adaptators was removed using Agencourt RNAClean XP beads. The RNA library was eluted from the RNA clean beads in 21 µl of elution buffer. 1 µl of the RNA library was quantified using a Qubit fluorometer using the manufacturer’s DNA HS assay. Immediately before sequencing, the remaining 20 µl of RNA library was mixed with 17.5 µl of nuclease-free water and 37.5 µl of RNA Running Buffer, making 75 µl of the final RNA library. The final RNA libraries were added to 9.4.1 flowcells (FLO-MIN106) and run on an Mk1b MinION.

The raw electrical signal from an ONT sequencer was translated to a RNA sequence using Oxford Nanopore basecaller, Albacore v2.3.3. The data quality was assessed using poretools v0.6.0 (1).

***RNA Sequencing with illumina sequencing system***

RNA-Seq libraries were constructed with the Truseq stranded mRNA sample preparation (Low throughput protocol) kit from illumina. One microgram of total RNA was used for the construction of the libraries.The first step in the workflow involves purifying the poly-A containing mRNA molecules using poly-T oligo attached magnetic beads. Following purification, the mRNA fragments are copied into first strand cDNA using SuperScript II reverse transcriptase, Actinomycine D and random hexamer primers. The Second strand cDNA was synthesized by replacing dTTP with dUTP. These cDNA fragments then have the addition of a single 'A' base and subsequent ligation of the adapter. The products are then purified and enriched with 15 cycles of PCR. The final cDNA libraries were validated with a Fragment Analyzer (Advanced Analytical, Ankeny, IA) and quantified with a KAPA qPCR kit (Roche-Kapa Biosystems, Wilmington, MA).

For each sequencing lane of a Rapid Run flowcell V2 or a High Throughput Run, 9 libraries were pooled in equal proportions, denatured with NaOH and diluted to 20 pM before clustering. Cluster formation, primer hybridisation and pair end-read 125 cycles sequencing were performed on cBot and HiSeq2500 (Illumina, San Diego, CA) respectively.

**MCF10A Cell culture**

Human mammary epithelial MCF10A-Snail-ER cells, which stably express the epithelial repressor Snaill fused to the estrogen receptor, were cultured as previously described in (2). Briefly, the cells were grown in DMEM/F-12 (Sigma) supplemented with 5% horse serum (Thermofisher), 20 ng/ml EGF (Sigma), 0.5 µg/ml hydrocortisone (Sigma), 100 ng/ml cholera toxin (Sigma), 10 µg/ml insulin (Sigma), 2mM L-Glutamine (Thermofisher) and 1x Pen./Strep. (Sigma). To induce epithelial-to-mesenchymal transition initiation, MCF10-Snail-ER cells were treated with 100 nM tamoxifen (Sigma) for 1 day. Importantly, parental MCF10A cells did not show any differences in cell morphology or transcriptional and splicing patterns upon the same treatment. All cell lines were regularly tested for mycoplasma presence.

**RNA extraction:**

After resuspending cell pellets in Trizol (Life technologies), total RNA was extracted by addition of chloroform and isopropanol precipitation, followed by RNA clean-up using the GeneJET RNA purification kit (Thermofisher).

**Long read data sets used for evaluation**

We adapted NanoSim (3) scripts so as to generate full-length ONT-like long reads mimicking the error profiles of our real data set. Our Long-read simulation procedure is set up as follows. ­­­­

Starting with a long-read data set, sequences are aligned on a reference transcriptome with Minimap2 (4). Then, NanoSim is used to estimate the parameters of an error-model for the long reads (mismatch, deletion and insertion rates and their length empirical distribution). Long reads are simulated according to this error-model and from a set of reference transcripts. The number of simulated LRs is proportional to transcript expression levels measured by real LRs. This is measured by following the method described in (5) where transcript expression levels are estimated using Salmon and annotation from ENSEMBL (release 97). We used this approach for two LR experiments from two distinct samples (our own MCF10A and downloaded GM12878 B-Lymphocyte cell lines) with two separate error models.

For the GM12878 cell line, the LR data was available from the Nanopore consortium at <https://github.com/nanopore-wgs-consortium/NA12878>. We made use of the *Run1* (MinION ONT direct-RNA, kit SQK-RNA001, pore R9.4) generated by the UCSC laboratory. These LRs were corrected using short read data from the same cell line sequenced by a separate consortium. These data were available from the GEO website ([https://www.ncbi.nlm.nih.gov/sra/SRX159827)](https://www.ncbi.nlm.nih.gov/sra/SRX15982)). After quality control using FastQC, we kept and pooled together runs SRR521447, SRR521448, SRR521453, SRR521454 and SRR521455. The discrepancy between the total numbers of real and simulated LRs in uneven experiments can be explained by the important part of truncated LRs whose isoform of origin remains ambiguous.

The SIRV (E0) long read dataset was downloaded at <https://www.ebi.ac.uk/ena/data/view/ERX3584788>.

These are cDNA reads generated on the Oxford Nanopore GridION system (pore R9) (6).

SIRV short reads used for correction were sequenced on Illumina HiSeq 2500. They are part of the experiments performed in (7), and were dowloaded at [https://www.ncbi.nlm.nih.gov/sra/SRX1756197[accn](https://www.ncbi.nlm.nih.gov/sra/SRX1756197%5Baccn)].

**Real Long Read datasets error rates estimates from NanoSim**

| **Model** | **Mismatch rate** | **Insertion rate** | **Deletion rate** | **Total error rate** |
| --- | --- | --- | --- | --- |
| **GM12878** | 0.040 | 0.033 | 0.078 | 0.15 |
| **MCF10A** | 0.058 | 0.035 | 0.111 | 0.20 |

**Total number of reads per dataset**

| **Model** | **# of Simulated LRs** | **# of Real LRs** | **# of Real SRs** |
| --- | --- | --- | --- |
| **SIRV** | - | 1 680 000 | 1 455 306 |
| **GM12878** | 162 373 | 488 754 | 105 171 449 |
| **MCF10A** | 104 101 | 319 146 | 104 819 581 |

**Estimated and** **Simulated transcript expression levels**

| **Model** | **Min.** | **1st quartile** | **Median** | **Mean** | **3rd quartile** | **Max.** |
| --- | --- | --- | --- | --- | --- | --- |
| **MCF10A** | 1 | 1 | 3 | 8,917 | 7 | 1 426 |
| **GM12878** | 1 | 2 | 4 | 37.98 | 16 | 16 907 |

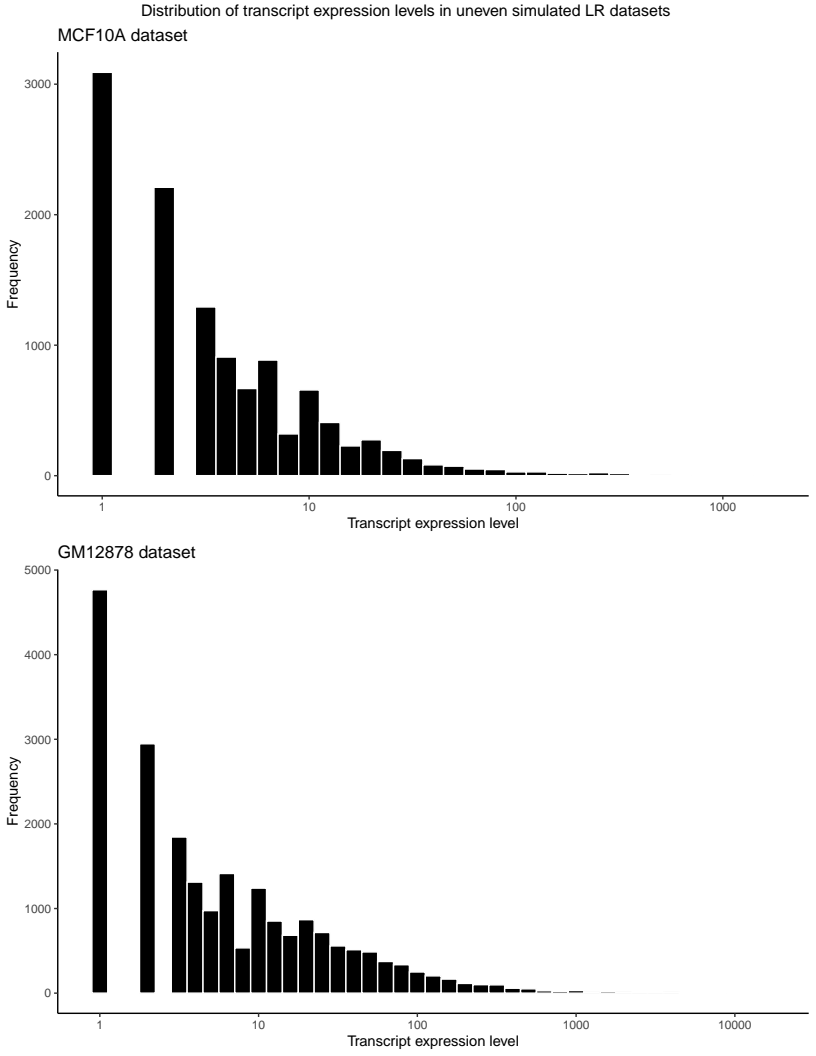

**Distribution of** **transcript expression levels in simulated datasets**

**Base accuracy computation**

Long reads were mapped to a reference genome with Minimap2, GMAP or to a reference transcriptome with GraphMap (and Minimap2 in the case of SIRV data). Then, only the primary alignment is considered and the base accuracy score is calculated as:

$baseaccuracy=\frac{nbM+nbIns+nbDel-NM}{nbM+nbIns+nbDel}$

where *nbM*, *nbIns* and *nbDel* are respectively the number of matching, inserted, and deleted bases and *NM* the edit distance reported by the aligner.

**Supplementary Table 1A: Usual alignment-based statistics for the 7 correction methods tested: primary alignment rates and base accuracy (real LR data corrected by real SR data).** Only the best reported alignment is counted. The two best performance scores are colored in green and the worst in red. Base accuracy is calculated as described above (cf: **Base accuracy computation**). Complementary summary statistics, error rates and similar results on simulated datasets can be found below in Supplementary Table 1B.

| **Alignments and Base Accuracy** | | | | | | | | | | | |
| --- | --- | --- | --- | --- | --- | --- | --- | --- | --- | --- | --- |
| **Method** | **GMAP** | | | | **Minimap2** | | | | **Graphmap** | | |
|  | **Percent primary alignments (%)** | **Mean Base accuracy** | **Median**  **Base accuracy** | **Percent primary alignments (%)** | | **Mean Base accuracy** | **Median**  **Base accuracy** | **Percent primary alignments (%)** | | **Mean Base accuracy** | **Median**  **Base accuracy** |
| **Real SIRV Spike-In data**  **(GridION cDNA)** | | | | | | | | | | | |
| **Raw** |  |  |  | 98.79 | | 92.28 | 92.96 | 99.01 | | 90.88 | 92.21 |
| **FMLRC** |  |  |  | 99.19 | | 94.56 | 99.46 | 99.25 | | 96.42 | 99.23 |
| **CoLoRMap** |  |  |  | 98.80 | | 92.64 | 93.39 | 21.73 | | 67.29 | 64.12 |
| **LoRDEC** |  |  |  | 99.12 | | 97.31 | 97.73 | 98.91 | | 94.60 | 96.74 |
| **TALC** |  |  |  | 99.08 | | 97.91 | 98.72 | 99.17 | | 96.70 | 98.33 |
| **Real MCF10A**  **(MinION direct-RNA)** | | | | | | | | | | | |
| **Raw** | 46.48 | 83.44 | 83.33 | 79.15 | | 81.16 | 81.55 | 93.04 | | 79.58 | 80.05 |
| **FMLRC** | 82.81 | 96.89 | 99.13 | 91.26 | | 94.07 | 97.34 | 91.74 | | 95.16 | 98.25 |
| **CoLoRMap** | 92.48 | 98.33 | 99.35 | 93.89 | | 97.45 | 99.26 | 83.62 | | 91.99 | 95.46 |
| **Hercules** | 80.88 | 90.93 | 91.57 | 91.19 | | 87.29 | 87.90 | 91.41 | | 84.96 | 85.65 |
| **LoRDEC** | 90.01 | 94.54 | 95.24 | 92.86 | | 92.35 | 93.70 | 82.27 | | 82.61 | 85.73 |
| **TALC** | 88.68 | 97.04 | 98.68 | 92.61 | | 94.97 | 97.73 | 93.81 | | 93.16 | 96.30 |
| **Real GM12878**  **(MinION direct-RNA)** | | | | | | | | | | | |
| **Raw** | 82.86 | 86.98 | 87.29 | 97.29 | | 86.04 | 86.63 | 96.40 | | 84.28 | 85.16 |
| **FMLRC** | 96.12 | 97.36 | 98.90 | 97.94 | | 96.35 | 98.17 | 96.89 | | 94.51 | 96.94 |
| **CoLoRMap** | 98.16 | 97.03 | 98.31 | 98.48 | | 96.19 | 98.29 | 82.72 | | 94.00 | 97.14 |
| **LoRDEC** | 98.48 | 96.12 | 97.03 | 98.16 | | 95.44 | 96.14 | 95.14 | | 90.44 | 93.51 |
| **TALC** | 97.98 | 97.41 | 98.32 | 98.11 | | 96.54 | 97.94 | 97.12 | | 94.96 | 96.86 |

**Supplementary Table 1B: General alignment statistics for the 7 correction methods tested on real LR datsets.** This table is similar to Supplementary Table 1A except that it includes minimum, first and third quantile measures in addition to median base accuracy (computed on all primary alignments).

|  | **Real SIRV Spike-In data** | | | | | | | | | | | |
| --- | --- | --- | --- | --- | --- | --- | --- | --- | --- | --- | --- | --- |
|  | **GraphMap** | | | | **GMAP** | | | | **Minimap2** | | | |
| **Base accuracy** | **Min.** | **1^st^ quant.** | **Median** | **3^rd^ quant.** | **Min.** | **1^st^ quant.** | **Median** | **3^rd^ quant.** | **Min.** | **1^st^ quant.** | **Median** | **3^rd^ quant.** |
| **Raw** | 55.25 | 88.87 | 92.21 | 94.33 |  |  |  |  | 45.75 | 90.42 | 92.96 | 94.79 |
| **FMLRC** | 55.08 | 96.91 | 99.23 | 100 |  |  |  |  | 43.59 | 95.74 | 99.46 | 100 |
| **CoLoRMap** | 20.17 | 53.92 | 64.12 | 78.97 |  |  |  |  | 42.59 | 90.68 | 93.39 | 95.51 |
| **LoRDEC** | 22.25 | 94.02 | 96.74 | 97.85 |  |  |  |  | 46.42 | 96.97 | 97.73 | 98.26 |
| **TALC** | 54.97 | 96.86 | 98.33 | 98.98 |  |  |  |  | 43.28 | 97.92 | 98.72 | 99.22 |

|  | **Real GM12878** | | | | | | | | | | | |
| --- | --- | --- | --- | --- | --- | --- | --- | --- | --- | --- | --- | --- |
|  | **GraphMap** | | | | **GMAP** | | | | **Minimap2** | | | |
| **Base accuracy** | **Min.** | **1^st^ quant.** | **Median** | **3^rd^ quant.** | **Min.** | **1^st^ quant.** | **Median** | **3^rd^ quant.** | **Min.** | **1^st^ quant.** | **Median** | **3^rd^ quant.** |
| **Raw** | 53.79 | 81.97 | 85.15 | 87.55 | 45.71 | 85.10 | 87.30 | 89.18 | 0.16 | 84.15 | 86.65 | 88.62 |
| **FMLRC** | 53.33 | 92.34 | 96.94 | 99.11 | 43.58 | 96.85 | 98.90 | 99.80 | 0.16 | 95.33 | 98.29 | 99.55 |
| **CoLoRMap** | 38.22 | 93.51 | 97.14 | 98.57 | 51.58 | 97.08 | 98.31 | 99.14 | 0.11 | 96.26 | 98.17 | 98.94 |
| **LoRDEC** | 35.76 | 87.13 | 93.51 | 96.92 | 50.00 | 94.67 | 96.40 | 98.45 | 0.16 | 94.06 | 96.15 | 98.13 |
| **TALC** | 53.82 | 93.44 | 96.86 | 98.63 | 58.53 | 96.59 | 98.32 | 99.28 | 0.16 | 95.71 | 97.94 | 99.11 |
|  | **Real MCF10A** | | | | | | | | | | | |
|  | **GraphMap** | | | | **GMAP** | | | | **Minimap2** | | | |
| **Base accuracy** | **Min.** | **1^st^ quant.** | **Median** | **3^rd^ quant.** | **Min.** | **1^st^ quant.** | **Median** | **3^rd^ quant.** | **Min.** | **1^st^ quant.** | **Median** | **3^rd^ quant.** |
| **Raw** | 53.71 | 77.11 | 80.05 | 82.55 | 49.03 | 81.15 | 83.33 | 85.52 | 0.14 | 79.15 | 81.55 | 83.71 |
| **FMLRC** | 53.77 | 86.54 | 95.46 | 99.26 | 33.33 | 96.11 | 99.13 | 100 | 0.14 | 90.69 | 97.34 | 99.76 |
| **CoLoRMap** | 27.70 | 95.28 | 98.25 | 99.37 | 38.27 | 98.78 | 99.35 | 99.75 | 0.40 | 98.29 | 99.26 | 99.70 |
| **Hercules** | 54.00 | 80.37 | 85.65 | 90.21 | 54.83 | 87.67 | 91.57 | 94.85 | 0.34 | 83.21 | 87.90 | 92.06 |
| **LoRDEC** | 25.07 | 74.12 | 85.73 | 93.48 | 47.78 | 92.65 | 95.24 | 97.02 | 0.12 | 89.87 | 93.70 | 96.08 |
| **TALC** | 53.97 | 89.92 | 96.30 | 98.99 | 42.78 | 96.03 | 98.68 | 99.71 | 0.14 | 92.78 | 97.73 | 99.46 |

**Supplementary Table 1C: Average Mismatch, Insertion and Deletion rates (real LR data corrected by real SR data)**

| **Error rates** | | | | | | | | | | |
| --- | --- | --- | --- | --- | --- | --- | --- | --- | --- | --- |
| **Method** | **GMAP** | | | **Minimap2** | | | **Graphmap** | | | |
|  | **%Mismatch** | **%Insertion** | **%Deletions** | **%Mismatch** | **%Insertion** | **%Deletions** | | **%Mismatch** | **%Insertion** | **%Deletion** |
| **Real SIRV Spike-Ins**  **(GridION cDNA)** | | | | | | | | | | |
| **Raw** |  |  |  | 2.505 | 2.230 | 2.984 | | 2.9345 | 2.835 | 3.353 |
| **FMLRC** |  |  |  | 0.226 | 1.122 | 4.087 | | 1.125 | 0.812 | 1.639 |
| **CoLoRMap** |  |  |  | 2.486 | 2.036 | 2.837 | | 31.318 | 0.561 | 0.827 |
| **LoRDEC** |  |  |  | 1.696 | 0.307 | 0.684 | | 4.3920 | 0.2613 | 0.748 |
| **TALC** |  |  |  | 0.949 | 0.294 | 0.848 | | 1.6000 | 0.831 | 0.872 |
| **Real MCF10A**  **(MinION direct-RNA)** | | | | | | | | | | |
| **Raw** | 4.442 | 5.143 | 6.967 | 5.38 | 3.302 | 10.148 | | 5.576 | 4.157 | 10.685 |
| **FMLRC** | 0.903 | 0.818 | 1.384 | 1.726 | 0.928 | 3.273 | | 2.390 | 1.560 | 4.062 |
| **CoLoRMap** | 0.613 | 0.521 | 0.530 | 1.047 | 0.367 | 1.135 | | 3.740 | 0.398 | 0.707 |
| **Hercules** | 1.011 | 3.892 | 4.160 | 2.976 | 3.029 | 6.698 | | 3.830 | 3.738 | 7.470 |
| **LoRDEC** | 2.232 | 1.453 | 1.770 | 3.238 | 0.994 | 3.412 | | 14.94 | 0.390 | 2.061 |
| **TALC** | 0.992 | 0.848 | 1.113 | 1.670 | 0.848 | 2.511 | | 2.250 | 1.389 | 3.199 |
| **Real GM12878**  **(MinION direct-RNA)** | | | | | | | | | | |
| **Raw** | 3.734 | 3.950 | 5.331 | 3.855 | 3.281 | 6.909 | | 4.202 | 4.146 | 7.37 |
| **FMLRC** | 0.882 | 0.653 | 1.102 | 1.134 | 0.736 | 1.939 | | 1.742 | 1.341 | 2.403 |
| **CoLoRMap** | 1.469 | 0.694 | 0.804 | 1.756 | 0.532 | 1.356 | | 5.046 | 0.358 | 0.592 |
| **LoRDEC** | 2.124 | 0.745 | 1.011 | 2.409 | 0.574 | 1.570 | | 7.832 | 0.493 | 1.232 |
| **TALC** | 1.048 | 0.728 | 0.816 | 1.252 | 0.749 | 1.458 | | 1.833 | 1.356 | 1.842 |

**Supplementary Tables 1D: General alignment statistics for the 7 correction methods tested on simulated LR datasets.** This table is similar to Supplementary Table 1A except that it includes first and third quantile measures in addition to median and mean base accuracy (computed on all primary alignments).

|  | **Simulated MCF10A** | | | | | | | | | | | | | | | | |
| --- | --- | --- | --- | --- | --- | --- | --- | --- | --- | --- | --- | --- | --- | --- | --- | --- | --- |
|  | **Graphmap** | | | | | | **GMAP** | | | | | | **Minimap2** | | | | |
| **Base accuracy** | **Min.** | **1^st^ quant.** | **Median** | **Mean** | **3^rd^ quant.** | **Min.** | | **1^st^ quant.** | **Median** | **Mean** | **3^rd^ quant.** | **Min.** | | **1^st^ quant.** | **Median** | **Mean** | **3^rd^ quant.** |
| **Raw** | 55.73 | 80.69 | 81.90 | 81.62 | 82.94 | 51.75 | | 81.95 | 83.22 | 83.47 | 84.74 | 0.40 | | 81.26 | 82.24 | 82.08 | 83.28 |
| **FMLRC** | 55.10 | 88.75 | 95.54 | 93.08 | 98.51 | 50.89 | | 93.71 | 97.70 | 95.70 | 99.63 | 0.69 | | 90.26 | 96.42 | 93.97 | 99.05 |
| **CoLoRMap** | 45.11 | 92.87 | 98.04 | 93.52 | 99.38 | 38.27 | | 98.78 | 99.35 | 98.33 | 99.75 | 0.76 | | 95.90 | 99.20 | 96.39 | 99.70 |
| **LSC** | 55.78 | 83.28 | 85.42 | 85.29 | 87.64 | 50.19 | | 85.88 | 88.19 | 88.54 | 91.01 | 0.35 | | 84.00 | 86.02 | 86.06 | 88.29 |
| **LoRDEC** | 30.25 | 72.95 | 82.93 | 80.70 | 90.26 | 38.14 | | 92.62 | 94.92 | 94.33 | 96.67 | 0.72 | | 91.19 | 93.92 | 93.14 | 96.03 |
| **TALC** | 55.11 | 91.90 | 96.08 | 94.39 | 98.85 | 52.01 | | 94.32 | 97.56 | 96.18 | 99.38 | 0.72 | | 92.68 | 96.69 | 95.07 | 99.16 |
|  | **Simulated GM12878** | | | | | | | | | | | | | | | | |
|  | **Graphmap** | | | | | | **GMAP** | | | | | | **Minimap2** | | | | |
| **Base accuracy** | **Min.** | **1^st^ quant.** | **Median** | **Mean** | **3^rd^ quant.** | **Min.** | | **1^st^ quant.** | **Median** | **Mean** | **3^rd^ quant.** | **Min.** | | **1^st^ quant.** | **Median** | **Mean** | **3^rd^ quant.** |
| **Raw** | 55.85 | 85.53 | 86.57 | 86.39 | 87.56 | 57.60 | | 85.97 | 87.06 | 87.16 | 88.33 | 0.42 | | 85.87 | 86.78 | 86.78 | 87.77 |
| **FMLRC** | 55.17 | 93.32 | 97.11 | 95.55 | 98.94 | 56.38 | | 94.72 | 97.99 | 96.51 | 99.43 | 0.11 | | 93.96 | 97.54 | 96.01 | 99.17 |
| **CoLoRMap** | 55.96 | 92.84 | 96.60 | 95.11 | 98.56 | 43.54 | | 95.05 | 98.13 | 96.38 | 99.11 | 0.24 | | 93.95 | 97.74 | 95.86 | 98.89 |
| **Hercules** | 55.30 | 90.85 | 92.56 | 92.08 | 94.0 | 57.94 | | 92.32 | 93.86 | 93.75 | 95.40 | 0.42 | | 91.68 | 93.17 | 93.01 | 94.61 |
| **LoRDEC** | 45.74 | 87.25 | 93.09 | 90.49 | 96.76 | 59.24 | | 94.98 | 96.61 | 96.40 | 98.44 | 0.48 | | 94.71 | 96.46 | 96.20 | 98.36 |
| **TALC** | 56.12 | 95.20 | 97.43 | 96.49 | 98.86 | 60.79 | | 95.87 | 97.90 | 97.02 | 99.11 | 0.66 | | 95.60 | 97.70 | 96.81 | 99.00 |

**Supplementary Table 1E:**

**Average base accuracy for the 7 correction methods tested relative to short read gene coverage.** This table is similar to Table 1 except that it displays average base accuracy relative to short read coverage of the reference gene to which the LR mapped. Reference genes were divided into three bins according to their number of short reads per base. Short reads were counted at the transcript-level using Salmon (5, 8) (quasi-mapping mode). Counts per transcript were then divided by the effective length of the transcript to estimate its “SR read depth”. The SR read depth of each gene was finally calculated by summing of the SR read depths over all its isoforms.

**SR gene coverage depth bins:**

| **Bins** | **Low** | **Medium** | **High** |
| --- | --- | --- | --- |
| **Short Read Depth ranges** | <1.5 | 1.5-25 | >25 |
| **GM12878 gene bin sizes** | 20 264 genes | 4 795 genes | 270 genes |
| **MCF10A gene bin sizes** | 14 185 genes | 4 547 genes | 416 genes |

|  | **Real GM12878** | | | | | | | | | | | |
| --- | --- | --- | --- | --- | --- | --- | --- | --- | --- | --- | --- | --- |
|  | **GraphMap** | | | | **GMAP** | | | | **Minimap2** | | | |
| **Base accuracy**  **per SR gene coverage bin** | **Zero** | **Low** | **Medium** | **High** | **Zero** | **Low** | **Medium** | **High** | **Zero** | **Low** | **Medium** | **High** |
| **Raw** | 83.47 | 83.85 | 84.11 | 84.61 | 87.06 | 86.32 | 86.86 | 87.57 | 82.66 | 85.12 | 85.65 | 86.17 |
| **FMLRC** | 92.71 | 94.02 | 95.92 | 93.87 | 96.30 | 96.30 | 98.24 | 97.24 | 89.93 | 96.58 | 96.89 | 95.51 |
| **CoLoRMap** | 93.09 | 94.18 | 94.27 | 93.81 | 96.82 | 96.82 | 97.31 | 96.95 | 89.90 | 96.58 | 96.57 | 96.01 |
| **LoRDEC** | 88.98 | 90.89 | 93.07 | 88.73 | 94.96 | 96.11 | 97.74 | 95.13 | 89.94 | 95.90 | 95.87 | 94.27 |
| **TALC** | 94.04 | 93.81 | 94.97 | 95.50 | 97.13 | 96.46 | 97.40 | 97.83 | 89.59 | 96.64 | 96.91 | 96.81 |
|  | **Real MCF10A** | | | | | | | | | | | |
|  | **GraphMap** | | | | **GMAP** | | | | **Minimap2** | | | |
| **Base accuracy**  **per SR gene coverage bin** | **Zero** | **Low** | **Medium** | **High** | **Zero** | **Low** | **Medium** | **High** | **Zero** | **Low** | **Medium** | **High** |
| **Raw** | 78.53 | 79.15 | 79.35 | 79.93 | 82.67 | 82.87 | 83.30 | 84.17 | 80.54 | 80.61 | 80.72 | 81.71 |
| **FMLRC** | 90.85 | 91.17 | 94.10 | 91.01 | 94.33 | 95.51 | 97.99 | 96.83 | 90.82 | 92.28 | 95.89 | 93.29 |
| **CoLoRMap** | 94.53 | 94.42 | 95.37 | 95.20 | 98.02 | 97.75 | 98.45 | 98.55 | 97.15 | 96.66 | 97.55 | 97.68 |
| **Hercules** | 82.74 | 83.51 | 84.39 | 86.06 | 90.09 | 89.29 | 90.06 | 92.36 | 85.91 | 85.31 | 86.24 | 88.73 |
| **LoRDEC** | 80.87 | 83.45 | 82.27 | 82.61 | 92.86 | 94.66 | 95.61 | 93.86 | 91.43 | 92.39 | 93.02 | 91.94 |
| **TALC** | 91.85 | 91.00 | 93.11 | 94.06 | 95.95 | 95.56 | 97.19 | 97.64 | 93.55 | 92.68 | 94.81 | 95.84 |

**Supplementary Table and Figure 1F: Relative distance between corrected and raw read length (real LR data sets).**

Relative distances were computed as : $\frac{Correctedreadlength-Rawreadlength}{Rawreadlength}$.

In all cases, the average and median distances are slightly over zero, consistent with Nanopore data higher deletion than insertion rate.

| **Absolute Relative distance between corrected and raw reads (x100)** | | | | | | | | | | | | | | | | | |
| --- | --- | --- | --- | --- | --- | --- | --- | --- | --- | --- | --- | --- | --- | --- | --- | --- | --- |
|  | **Real SIRV data** | | | | | | **Real MCF10A data** | | | | | | **Real GM12878 data** | | | | |
|  | **1^st^ quant.** | **Median** | **Mean** | **3^rd^ quant.** | **sd.** | **1^st^ quant.** | | **Median** | **Median** | **3^rd^ quant.** | **sd.** | **1^st^ quant.** | | **Median** | **Mean** | **3^rd^ quant.** | **sd.** |
| **FMLRC** | 0.699 | 1.623 | 2.175 | 3.079 | 2.04 | 0.79 | | 3.53 | 3.84 | 5.95 | 3.34 | 0.88 | | 2.14 | 2.57 | 3.68 | 2.39 |
| **CoLoRMap** | 0 | 0.133 | 0.607 | 0.716 | 1.27 | 7.66 | | 12.15 | 13.61 | 17.90 | 9.92 | 2.35 | | 4.45 | 5.02 | 6.82 | 4.07 |
| **Hercules** | More than a week | | | | | 0.94 | | 2.75 | 3.39 | 4.88 | 3.35 | More than a week | | | | | |
| **LoRDEC** | 0.316 | 0.712 | 1.475 | 1.295 | 7.28 | 2.39 | | 4.44 | 4.58 | 6.48 | 2.95 | 1.32 | | 2.60 | 2.93 | 4.11 | 2.10 |
| **TALC** | 0.364 | 0.808 | 1.098 | 1.460 | 1.18 | 2.51 | | 5.10 | 5.29 | 7.59 | 3.56 | 1.46 | | 2.79 | 3.15 | 4.36 | 2.27 |

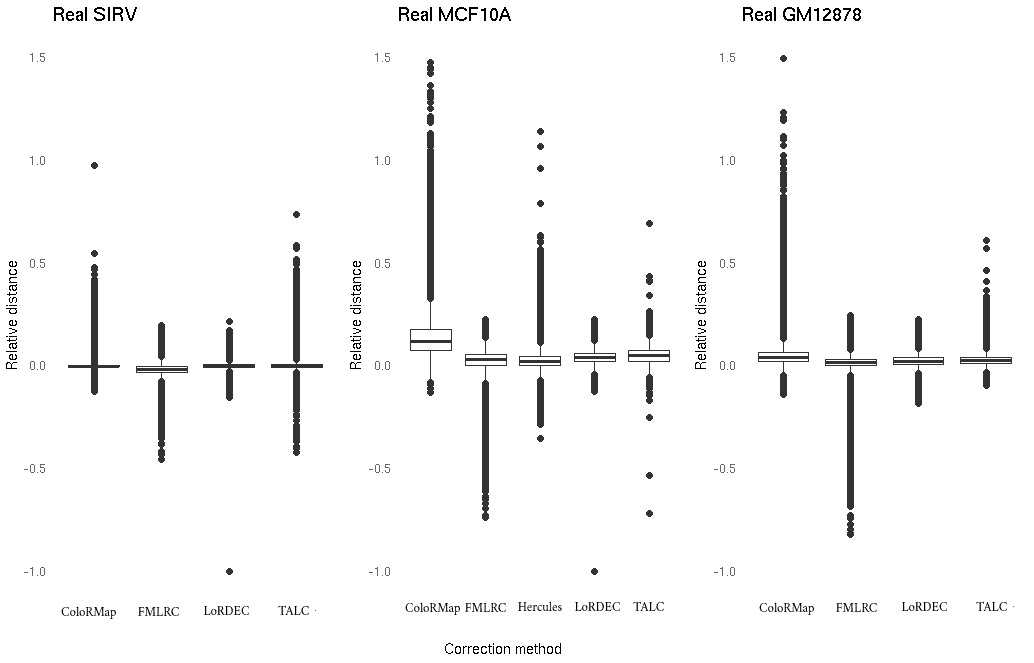

**Supplementary Table 2A: summaries of error rates and absolute relative distance** **in Salmon gene-level quantification compared to (known) ground truth (simulated LR data corrected by real SR data).** Gene expression levels were estimated by summing the Number of Reads obtained from various the transcript-level quantification strategies and their accuracy was evaluated using two measures: the absolute distance and the absolute relative distance (9) between the estimates and the ground truth.

|  |  | **Simulated MCF10A** | | | | | | | | |
| --- | --- | --- | --- | --- | --- | --- | --- | --- | --- | --- |
|  |  | **Absolute Error (gene-level quantification)** | | | | | | | | |
|  |  | **Salmon quasi-mapping** | **Minimap2+Salmon** | **Minimap2+Salmon** | | | | | | |
| **Method** | **1^st^ quant.** | **Median** | **Mean** | **3^rd^ quant.** | **sd** | **1^st^ quant.** | **Median** | **Mean** | **3^rd^ quant.** | **sd** |
| **Raw** | 2 | 4 | 12.38 | 9 | 49.14 | 1 | 2 | 8.90 | 5 | 45.93 |
| **FMLRC** | 1 | 2 | 6.30 | 4 | 29.31 | 1 | 2 | 8.24 | 5 | 39.43 |
| **CoLoRMap** | 1 | 2 | 6.77 | 5 | 28.64 | 1 | 2 | 7.11 | 4 | 34.88 |
| **LSC** | 1 | 3 | 10.29 | 8 | 35.82 | 0.5 | 2 | 8.42 | 42.99 | 42.99 |
| **LoRDEC** | 1 | 1 | 3.97 | 3 | 14.28 | 0 | 1 | 8.42 | 4 | 42.50 |
| **TALC** | 1 | 2 | 5.17 | 4 | 21.84 | 0 | 1 | 7.69 | 4 | 38.85 |
|  |  | **Absolute Error (gene-level quantification)** | | | | | | | | |
|  |  | **Salmon quasi-mapping** | **Minimap2+Salmon** | **Minimap2+Salmon** | | | | | | |
| **Method** | **1^st^ quant.** | **Median** | **Mean** | **3^rd^ quant.** | **sd** | **1^st^ quant.** | **Median** | **Mean** | **3^rd^ quant.** | **sd** |
| **Raw** | 1 | 1 | 0.99 | 1 | 0.05 | 0.07 | 1 | 0.67 | 1 | 0.45 |
| **FMLRC** | 0.29 | 1 | 0.69 | 1 | 0.40 | 0.14 | 1 | 0.69 | 1 | 0.44 |
| **CoLoRMap** | 0.25 | 1 | 0.68 | 1 | 0.41 | 0.06 | 1 | 0.67 | 1 | 0.45 |
| **LSC** | 1 | 1 | 0.91 | 1 | 0.22 | 0.01 | 1 | 0.66 | 1 | 0.46 |
| **LoRDEC** | 0.07 | 0.31 | 0.46 | 1 | 0.42 | 0.05 | 1 | 0.63 | 1 | 0.46 |
| **TALC** | 0.18 | 0.90 | 0.61 | 1 | 0.41 | 0 | 1 | 0.63 | 1 | 0.46 |
|  |  | **Simulated GM12878** | | | | | | | | |
|  |  | **Absolute Error (gene-level quantification)** | | | | | | | | |
|  |  | **Salmon quasi-mapping** | **Minimap2+Salmon** | **Minimap2+Salmon** | | | | | | |
| **Method** | **1^st^ quant.** | **Median** | **Mean** | **3^rd^ quant.** | **sd** | **1^st^ quant.** | **Median** | **Mean** | **3^rd^ quant.** | **sd** |
| **Raw** | 1 | 3 | 16.54 | 9 | 84.15 | 0 | 1 | 14.54 | 4 | 113.66 |
| **FMLRC** | 0 | 1 | 4.95 | 2 | 30.35 | 0 | 1 | 13.04 | 4 | 105.77 |
| **CoLoRMap** | 0 | 1 | 6.70 | 3 | 41.12 | 0 | 1 | 12.67 | 3 | 108.30 |
| **Hercules** | 0 | 0 | 2.68 | 1 | 14.73 | 0 | 1 | 12.27 | 3 | 103.28 |
| **LoRDEC** | 0 | 0 | 2.11 | 1 | 12.13 | 0 | 1 | 11.85 | 3 | 94.80 |
| **TALC** | 0 | 0.98 | 3.57 | 1.46 | 22.61 | 0 | 1 | 11.16 | 3 | 89.40 |
|  |  | **Absolute Error (gene-level quantification)** | | | | | | | | |
|  |  | **Salmon quasi-mapping** | **Minimap2+Salmon** | **Minimap2+Salmon** | | | | | | |
|  | **1^st^ quant.** | **Median** | **Mean** | **3^rd^ quant.** | **sd** | **1^st^ quant.** | **Median** | **Mean** | **3^rd^ quant.** | **sd** |
| **Raw** | 0.71 | 1 | 0.83 | 1 | 0.27 | 0 | 1 | 0.53 | 1 | 0.48 |
| **FMLRC** | 0 | 0.12 | 0.40 | 1 | 0.45 | 0 | 1 | 0.57 | 1 | 0.47 |
| **CoLoRMap** | 0 | 2 | 0.44 | 1 | 0.45 | 0 | 1 | 0.53 | 1 | 0.48 |
| **Hercules** | 0 | 0 | 0.23 | 0.24 | 0.39 | 0 | 0.33 | 0.50 | 1 | 0.48 |
| **LoRDEC** | 0 | 0 | 0.25 | 0.33 | 0.41 | 0 | 0.14 | 0.46 | 1 | 0.48 |
| **TALC** | 0 | 0 | 0.32 | 1 | 0.434 | 0 | 0.33 | 0.50 | 1 | 0.48 |

**Supplementary Table 2B: summaries of error rates and ARD in transcript-level quantification compared to (known) ground truth (simulated LR data corrected by real SR data).** As for Supplementary Table 2A**, t**he accuracy was evaluated using two measures: the absolute distance and the absolute relative distance (ARD) between the estimated read counts and the ground truth.

|  | **Simulated MCF10A** | | | | | | | | | | | | | | | | |
| --- | --- | --- | --- | --- | --- | --- | --- | --- | --- | --- | --- | --- | --- | --- | --- | --- | --- |
|  | **Absolute Error (transcript-level quantification)** | | | | | | | | | | | | | | | | |
|  | **Salmon quasi-mapping** | | | | | **Minimap2+Salmon** | | | | | | **GraphMap** | | | | | |
| **Method** | **1^st^ quant.** | **Median** | **Mean** | **3^rd^ quant.** | **sd** | | **1^st^ quant.** | **Median** | **Mean** | **3^rd^ quant.** | **sd** | | **1^st^ quant.** | **Median** | **Mean** | **3^rd^ quant.** | **sd** |
| **Raw** | 1 | 3 | 8.88 | 7 | 32.93 | | 0 | 1.45 | 7.87 | 4 | 30.93 | | 0.05 | 0.19 | 0.49 | 1 | 0.46 |
| **FMLRC** | 1 | 2 | 5.61 | 4 | 23.09 | | 1 | 2 | 7.58 | 5 | 29.7 | | 0.07 | 1 | 0.62 | 1 | 0.44 |
| **CoLoRMap** | 1 | 2 | 6.39 | 4 | 25.68 | | 0 | 1 | 6.78 | 4 | 30.16 | | 0.17 | 1 | 0.68 | 1 | 0.41 |
| **LSC** | 1 | 3 | 7.46 | 6 | 24.72 | | 0 | 1 | 7.36 | 4 | 29.61 | | 0.05 | 0.19 | 0.49 | 1 | 0.46 |
| **LoRDEC** | 0 | 1 | 3.20 | 2 | 11.69 | | 0 | 1 | 7.21 | 4 | 30.24 | | 0.09 | 0.24 | 0.51 | 1 | 0.45 |
| **TALC** | 1 | 1 | 4.56 | 3 | 18.68 | | 0 | 1 | 6.79 | 4 | 29.91 | | 0.06 | 0.34 | 0.52 | 1 | 0.45 |
|  | **Absolute Relative Distances** | | | | | | | | | | | | | | | | |
|  | **Salmon quasi-mapping** | | | | | **Minimap2+Salmon** | | | | | | **GraphMap** | | | | | |
| **Method** | **1^st^ quant.** | **Median** | **Mean** | **3^rd^ quant.** | **sd** | | **1^st^ quant.** | **Median** | **Mean** | **3^rd^ quant.** | **sd** | | **1^st^ quant.** | **Median** | **Mean** | **3^rd^ quant.** | **sd** |
| **Raw** | 1 | 1 | 0.99 | 1 | 0.048 | | 0 | 1 | 0.63 | 1 | 0.46 | | 0.05 | 0.19 | 0.49 | 1 | 0.46 |
| **FMLRC** | 0.29 | 1 | 0.69 | 1 | 0.40 | | 0.11 | 1 | 0.66 | 1 | 0.44 | | 0.07 | 1 | 0.62 | 1 | 0.44 |
| **CoLoRMap** | 0.21 | 1 | 0.66 | 1 | 0.41 | | 0 | 1 | 0.63 | 1 | 0.45 | | 0.16 | 1 | 0.69 | 1 | 0.41 |
| **LSC** | 1 | 1 | 0.89 | 1 | 0.22 | | 0 | 1 | 0.60 | 1 | 0.46 | | 0.04 | 0.19 | 0.49 | 1 | 0.46 |
| **LoRDEC** | 0 | 0.25 | 0.43 | 1 | 0.42 | | 0 | 1 | 0.57 | 1 | 0.47 | | 0.09 | 0.24 | 0.51 | 1 | 0.45 |
| **TALC** | 0.14 | 0.66 | 0.58 | 1 | 0.42 | | 0 | 1 | 0.57 | 1 | 0.46 | | 0.06 | 0.35 | 0.52 | 1 | 0.45 |
|  | **Simulated GM12878** | | | | | | | | | | | | | | | | |
|  | **Absolute Error (transcript-level quantification)** | | | | | | | | | | | | | | | | |
|  | **Salmon quasi-mapping** | | | | | **Minimap2+Salmon** | | | | | | **GraphMap** | | | | | |
| **Method** | **1^st^ quant.** | **Median** | **Mean** | **3^rd^ quant.** | **sd** | | **1^st^ quant.** | **Median** | **Mean** | **3^rd^ quant.** | **sd** | | **1^st^ quant.** | **Median** | **Mean** | **3^rd^ quant.** | **sd** |
| **Raw** | 1 | 3 | 11.38 | 7 | 57.73 | | 0 | 1 | 11.77 | 3 | 70.45 | | 0.28 | 1 | 4.02 | 2 | 39.53 |
| **FMLRC** | 0 | 1 | 4.96 | 2 | 30.01 | | 0 | 1 | 11.05 | 3 | 68.67 | | 0.61 | 1 | 5.56 | 3.42 | 30.64 |
| **CoLoRMap** | 0 | 1 | 6.36 | 3 | 39.43 | | 0 | 1 | 10.52 | 3 | 72.23 | | 1 | 1.87 | 7.24 | 2.16 | 39.53 |
| **Hercules** | 0 | 0 | 2.47 | 1 | 16.08 | | 0 | 1 | 9.76 | 3 | 66.33 | | 0.41 | 1 | 4.59 | 2.16 | 25.60 |
| **LoRDEC** | 0 | 0 | 2.26 | 1 | 14.20 | | 0 | 0 | 9.28 | 2 | 62.52 | | 0.31 | 1 | 3.47 | 1.71 | 19.64 |
| **TALC** | 0 | 0 | 3.72 | 1.16 | 26.09 | | 0 | 1 | 9.03 | 2.312 | 62.18 | | 0.32 | 1 | 4.37 | 1.86 | 29.23 |
|  | **Absolute Relative Distances** | | | | | | | | | | | | | | | | |
| **Method** | **Salmon quasi-mapping** | | | | | **Minimap2+Salmon** | | | | | | **GraphMap** | | | | | |
|  | **1^st^ quant.** | **Median** | **Mean** | **3^rd^ quant.** | **sd** | | **1^st^ quant.** | **Median** | **Mean** | **3^rd^ quant.** | **sd** | | **1^st^ quant.** | **Median** | **Mean** | **3^rd^ quant.** | **sd** |
| **Raw** | 0.67 | 1 | 0.83 | 1 | 0.28 | | 0 | 0.23 | 0.47 | 1 | 0.47 | | 0.03 | 0.11 | 0.46 | 1 | 0.47 |
| **FMLRC** | 0 | 0.15 | 0.41 | 1 | 0.45 | | 0 | 0.5 | 0.52 | 1 | 0.47 | | 0.05 | 0.43 | 0.52 | 1 | 0.46 |
| **CoLoRMap** | 0 | 0.18 | 0.41 | 1 | 0.45 | | 0 | 0.2 | 0.46 | 1 | 0.47 | | 0.18 | 1 | 0.61 | 1 | 0.43 |
| **Hercules** | 0 | 0 | 0.21 | 0.20 | 0.36 | | 0 | 0.08 | 0.43 | 1 | 0.47 | | 0.04 | 0.16 | 0.48 | 1 | 0.46 |
| **LoRDEC** | 0 | 0 | 0.22 | 0.26 | 0.39 | | 0 | 0 | 0.39 | 1 | 0.47 | | 0.04 | 0.04 | 0.44 | 1 | 0.46 |
| **TALC** | 0 | 0 | 0.29 | 0.71 | 0.42 | | 0 | 0.09 | 0.43 | 1 | 0.47 | | 0.04 | 0.05 | 0.44 | 1 | 0.46 |

**Supplementary Table 2C: Overall impact of hybrid correction methods on transcript structure (real SR data, simulated LR data).** For each aligner, we consider all exons from the transcripts that generated the LR if the LR mapped to the correct gene (#Exons evaluated). We computed the proportion of exons that were correctly mapped (%Identified). It can be interpreted as the exon recall. We then computed the same statistic for terminal and internal exons separately. We also report the fraction of full-length LRs for which all exons could be accurately identified (*error-free* structure). The set of exons that could be evaluated varies between tools as it depends on the reads that could be mapped to the correct gene of origin. An exon is considered as identified if its genomic position is fully overlapped and the exon boundaries recognized by the aligner differ by no more than 10 bp. False exons are estimated by measuring sequences longer than 25 bp introduced in the intronic region of the transcript of origin.

|  | **Simulated MCF10A transcriptome** | | | | | | | | | |
| --- | --- | --- | --- | --- | --- | --- | --- | --- | --- | --- |
| **Method** | **GraphMap** | | | **GMAP** | | | **Minimap2** | | | |
|  | **#Exons evaluated %Identified**  **(border, internal)** | **Average # of false exons per 100 LRs** | **%evaluated LR with error-free structure** | **#Exons evaluated %Identified**  **(border, internal)** | **Average # of false exons per 100 LRs** | **%evaluated LR with error-free structure** | | **#Exons evaluated %Identified**  **(border, internal)** | **Average # of false exons per 100 LRs** | **%evaluated LR with error-free structure** |
| **Raw** | 784,270 84.4  (83.7, 87.1) | 2.82 | 90.4 | 687,948 45.5  (40.6, 42.5) | 10 | 73.00 | | 768,693 82.7  (80.1, 86.8) | 8.1 | 72.3 |
| **FMLRC** | 784,201 86.5  (85.7, 94.7) | **9.24** | **82.1** | 733,559 76.6  (70.90, 81.7) | **14.26** | **70.3** | | 761,309 83.7  (80.1, 90.3) | **13.8** | **69.6** |
| **CoLoRMap** | 615,660 **89.8**  (88.5, 96.8) | 7.7 | 85.4 | 760,483 **81.4**  (77.5, 85.9) | 8.69 | 78.6 | | 775784 **88.7**  (86.4, 94.5) | 10.67 | **78.4** |
| **LSC** | 782,747 86.2  (85.8, 89.8) | **3.17** | **90.7** | 730,408 **52.5**  (48.5, 52.1) | 8.54 | 74.9 | | 775,160 83.1  (81.1, 88.2) | **7.3** | 74.8 |
| **LoRDEC** | 754,764 **84.8**  (84.0, 95.0) | **2.67** | **90.3** | 763,388 77.9  (74.3, 81.5) | **3.47** | **82.2** | | 777,363 **87.4**  **(85.1, 93.1)** | **4.36** | 78.3 |
| **TALC** | 784,471 **89.7**  (88.7, 96.5) | 5.23 | 90.2 | 762,436 **80.8**  (77.1,86.8) | **6.68** | **82.9** | | 774,031 **87.4**  (85, 94.4) | 7.5 | **81.4** |
|  | **Simulated GM12878 transcriptome** | | | | | | | | | |
| **Raw** | 1,146,151 88.9  (88.2, 90.8) | 1.47 | 94.6 | 1,091,800 63.4  (59.7, 65.4) | 3.99 | 86.9 | | 1,124,919 89  (86.7, 92.5) | 3.49 | 84.1 |
| **FMLRC** | 1,146,613 91  (90.4, 96.7) | **3.83** | **91.5** | 1,108,002 85.5  (81.5, 89.6) | 5.27 | **85.8** | | 1,112,787 88.9  (85.6, 93.1) | **5.54** | **82.3** |
| **CoLoRMap** | 1,144,432 **92.9**  **(92.0, 96.5)** | 2.90 | 93.6 | 1,054,935 84.1  (76.1, 85.7) | **5.39** | 88 | | 1,119,129 **90.3**  (87.5, 94.8) | 5.28 | 85.9 |
| **Hercules** | 1,142,374 87.8  (87.1, 92.9) | 2.72 | 92.6 | 1,128,603 **73.6**  **(**71.5, 78.5) | **1.93** | 90.8 | | 1,128,585 85.9  (83.9, 91.7) | **2.65** | 87.7 |
| **LoRDEC** | 1,140,409 89.2  (88.6, 97.7) | **1.24** | **95.8** | 1,138,913 **90.2**  (88.4, 94.1) | **0.91** | **93.9** | | 1,124,919 89  (87.7, 94.3) | **1.54** | **90.1** |
| **TALC** | 1,147,448 **92.6**  (91.9, 98.1) | **2.40** | **95.5** | 1,123,019 87.8  (84.8, 92.9) | 2.46 | **93.3** | | 1,122,112 **91.1**  (88.5, 96.2) | 2.78 | **91.6** |

**Supplementary** **Table 2D: Improvement of hybrid correction methods on local transcript structure compared to raw data (real SR data, simulated LR data).** To evaluate the improvement on exon identification, we computed the number of exons properly mapped in raw data that are still properly mapped after correction (number of exons preserved) and the number of exons missed by the aligner in the raw data that are properly assigned after correction (number of exons clarified). For each correction method, we only consider LRs that aligned to the correct genomic locus both in the raw and in the corrected data set.

| **Simulated MCF10A transcriptome** | | | | | | | | |
| --- | --- | --- | --- | --- | --- | --- | --- | --- |
| **Method** | **GraphMap** | | **GMAP** | | | **Minimap2** | | |
|  | **#Raw True #Preserved %Preserved** | **#Raw False #Clarified %Clarified** | | **#Raw True #Preserved %Preserved** | **#Raw False #Clarified %Clarified** | | **#Raw True #Preserved %Preserved** | **#Raw False #Clarified %Clarified** |
| **FMLRC** | 655610 633408 96.6% | 10070 4854 48.2% | | 297882 279929 **94%** | 26839 13434 50% | | 617044 591339 **95.8%** | 34182 13871 40.6% |
| **CoLoRMap** | 511900 496570 97% | 7838 5027 **64%** | | 302854 287853 95% | 27471 15789 **57.5%** | | 626080 606432 96.9% | 34597 20573 **59.5%** |
| **LSC** | 657401 654678 **99.6%** | 10030 2107 **21%** | | 305775 290263 94.9% | 27226 6248 **22.9%** | | 630779 626339 **99.3%** | 34638 5111 **14.8%** |
| **LoRDEC** | 632786 608172 **96.1%** | 9769 5695 58.3% | | 306702 301385 **98.3%** | 27320 13652 50% | | 628474 624944 **99.4%** | 34550 12556 36% |
| **TALC** | 655488 647047 **98.7%** | 10051 6588 **65.5%** | | 303886 296934 **97.7%** | 27367 16731 **61.1%** | | 624881 615650 98.5% | 34452 19512 **56.6%** |
| **Simulated GM12878 transcriptome** | | | | | | | | |
| **FMLRC** | 1012185 991313 97.9% | 8519 4840 56.8% | | 669328 646408 96.6% | 18031 8565 47.5% | | 977535 953327 97.5% | 27356 9159 33.5% |
| **CoLoRMap** | 1011390 998538 **98.7%** | 8549 4914 57.5% | | 639460 61722 96.5% | 17146 10018 58.4% | | 983379 959073 97.5% | 27093 13154 **48.6%** |
| **Hercules** | 101033 992817 98.2% | 8493 2772 **32.6%** | | 681744 640724 **94%** | 18107 8123 **44.9%** | | 989334 965503 97.6% | 27592 8937 **32.4%** |
| **LoRDEC** | 1006423 984650 **97.8%** | 8499 5603 **65.9%** | | 684944 678472 **99.1%** | 18416 11591 **62.9%** | | 989733 985834 **99.6%** | 27657 13055 47.2% |
| **TALC** | 1012506 1003879 **99.1%** | 8600 6118 **71.1%** | | 674515 666202 **98.8%** | 18273 11949 **65.4%** | | 983063 974064 **99.1%** | 27494 16366 **59.5%** |

**Supplementary Table 2E: Impact of hybrid correction methods on Spike-In transcript quantification (real LR data corrected by real SR data).** Transcript expression levels were computed using two approaches: the first is based on Salmon (quasi-mapping mode)  and the other one on transcriptome assignments using GraphMap. The expected number of reads per Spike-In transcript was then estimated by multiplying theoretical concentrations by the size of the library (1 680 000 LRs). Error rates were finally obtained as the absolute value of the difference: *Expected value – Estimated value*. To account for diverging transcript expression levels we also reported relative errors (*Expected value – Estimate)/Expected value.*

|  | **Real SIRV data** | | | | | | | | | | |
| --- | --- | --- | --- | --- | --- | --- | --- | --- | --- | --- | --- |
|  | **Absolute Errors** | | | | | | | | | | |
|  | **Minimap2+Salmon**  **(transcriptome alignment)** | | | | | **Graphmap**  **(transcriptome alignment)** | | | | | |
| **Method** | **1^st^ quant.** | **Median** | **Mean** | **3^rd^ quant.** | **sd** | | **1^st^ quant.** | **Median** | **Mean** | **3^rd^ quant.** | **sd** |
| **Raw** | 7163 | 19455 | 28770 | 36406 | 33346 | | 9484 | 19562 | 31445 | 39469 | 35800 |
| **FMLRC** | 9015 | 17460 | 23722 | 30398 | 24618 | | 7591 | 18223 | 29247 | 40632 | 35534 |
| **CoLoRMap** | 7569 | 18587 | 27838 | 36279 | 32557 | | 10268 | 20230 | 25135 | 36878 | 19726 |
| **LoRDEC** | 8065 | 19447 | 28746 | 36924 | 33242 | | 9350 | 20073 | 31646 | 39616 | 35965 |
| **TALC** | 6576 | 19235 | 27464 | 36161 | 31357 | | 9221 | 19949 | 31261 | 39653 | 35761 |
|  | **Relative Errors** | | | | | | | | | | |
|  | **Minimap2+Salmon**  **(transcriptome alignment)** | | | | | **Graphmap**  **(transcriptome alignment)** | | | | | |
| **Method** | **1^st^ quant.** | **Median** | **Mean** | **3^rd^ quant.** | **sd** | | **1^st^ quant.** | **Median** | **Mean** | **3^rd^ quant.** | **sd** |
| **Raw** | 0.292 | 0.495 | 0.513 | 0.732 | 0.269 | | 0.310 | 0.577 | 0.559 | 0.813 | 0.287 |
| **FMLRC** | 0.305 | 0.466 | 0.494 | 0.700 | 0.275 | | 0.233 | 0.531 | 0.527 | 0.802 | 0.308 |
| **CoLoRMap** | 0.288 | 0.472 | 0.491 | 0.721 | 0.260 | | 0.271 | 0.464 | 0.466 | 0.6409 | 0.256 |
| **LoRDEC** | 0.288 | 0.498 | 0.512 | 0.729 | 0.267 | | 0.304 | 0.582 | 0.565 | 0.817 | 0.287 |
| **TALC** | 0.291 | 0.488 | 0.495 | 0.697 | 0.259 | | 0.294 | 0.566 | 0.559 | 0.807 | 0.290 |

**Supplementary Table 3: (real LR data corrected by real SR data) Runtimes and memory consumption on the real datasets.**

All methods were run on 15 CPUs.

|  | **SIRV** | | **MCF10A** | | | **GM12878** | | |
| --- | --- | --- | --- | --- | --- | --- | --- | --- |
| **Method** | **Runtime**  **(h:mm:ss)** | **Memory Peak (GB)** | **Runtime**  **(h:mm:ss)** | | **Memory Peak (GB)** | | **Runtime**  **(h:mm:ss)** | **Memory Peak (GB)** |
| **FMLRC** | **00:19:26** | **0.45** | **00:31:30** | | **16.8** | | **00:33:00** | **21.6** |
| **CoLoRMap** | **01:49:22** | **51.4** | **~2 days** | | **20** | | **12:41:45** | **21.15** |
| **Hercules** | **More than a week** | | **~6 days** | **10** | | **More than a week** | | |
| **LoRDEC** | **02:42:29** | **0.44** | **00:59:05** | | **2.65** | | **02:08:54** | **2.27** |
| **TALC** | **06:44:45** | **3.5** | **04:29:45** | | **34** | | **03:50:52** | **42** |

**Modeling of k-mer counts and node classification**

When considering a specific node n0 and one of its successors n1, there are three possible configurations:

1) n0 and n1 weights are not statistically different;

2) there is a significant difference in coverage between n1 and n0 and the n1/n0 fraction is close to the error rate expected in SR;

3) there is a significant difference in coverage between n1 and n0 but the n1/n0 fraction is higher than the error rate expected in SR;

In other words, each node can be classified, knowing its predecessor, either as *count-consistent*, *change-point* or *background noise*.

The above behavior of k-mer counts (or node weights) is modeled through Poisson distributions, and nodes are tagged using the following rational:

1) Knowing n0, if n1 is consistent with n0, we expect n1 to fluctuate around n0; which is formalized as: n1 ~ Poisson(n0) (H0).

This hypothesis is tested against its alternative n1 ~ Poisson(k) k != n0 (H1).

If H0 is not rejected, n1 is considered consistent with n0.

If H0 is rejected, it may be that n1 is noise. Then we expect that n1 ~ Poisson(mu) where mu is less than the average background noise around n0, which we estimate as lambda=n0*epsilon where epsilon is the per base error rate in SR).

This hypothesis is tested against (H1) n1~Poisson(mu), mu>lambda.

If it is rejected as well, n1 is classified as a change-point in the path, otherwise the k-mer is assumed to stem from a sequencing error.

**Confidence intervals to detect inconsistent k-mer counts**

In TALC, abrupt count variations are defined as counts that depart from what is expected by the Poisson model of the current k-mer count. More precisely, we compute a Poisson confidence interval around the current k-mer count (see below) and consider that a k-mer is not count-consistent if its count falls outside this interval.

In addition, to test whether a k-mer is likely erroneous because it’s counts are below noise levels, we compute a Poisson confidence interval around the expected noise level (**cf: Modeling of k-mer counts and node classification**). If the k-mer counts is less than the upper bound of this interval, it is considered as noise.

To select a suitable method to compute Poisson confidence intervals we use Wald’s CC method if the k-mer counts are less than 4 (10) and Bégaud’s method (11) otherwise.

**Border refinement procedure**

When the elected path is longer than the reference, successive seed-and-extend alignments are performed with a decreasing number of errors until an optimal alignment position of the LR onto the path has been found. The path is then trimmed at that position.

If no long border path could be found, shorter paths are compared and the one with highest similarity to the LR is selected. Its alignment position onto the LR is found as described above by successive seed-and-extend alignments. The suffix of LR’s sequence starting at that position is then appended to the short path’s.
